# Evolutionary origin and functional mechanism of Lhcx in the diatom photoprotection

**DOI:** 10.1101/2025.09.06.674587

**Authors:** Minoru Kumazawa, Seiji Akimoto, Atsushi Takabayashi, Ko Imaizumi, Shoko Tsuji, Hazuki Hasegawa, Atsushi Sakurai, Sousuke Imamura, Noriko Ishikawa, Natsuko Inoue-Kashino, Yasuhiro Kashino, Kentaro Ifuku

## Abstract

Diatoms are red-lineage algae that utilize the light-harvesting complex (LHC) subfamily Lhcx for photoprotection via non-photochemical quenching (NPQ); however, its evolutionary origin and molecular mechanism remain poorly understood. Through molecular phylogenetic analysis, we show that diatom Lhcxs and green algal Lhcsrs evolved from a common ancestor, with green plants subsequently acquiring them via horizontal gene transfer. To investigate the functional role of Lhcx1, we generated knockout mutants of *Chaetoceros gracilis*, a diatom with low Lhcx redundancy. The *lhcx1* mutants nearly abolished NPQ, and time-resolved fluorescence measurements revealed that Lhcx1-mediated quenching occurs in energetically detached antenna complexes. Clear-native PAGE with Amphipol further indicated that CgLhcx1 interacts with the FCP L-dimer, functioning as a peripheral antenna for the C_2_S_2_M_2_ PSII–FCPII supercomplex. Notably, under high-light acclimation, *lhcx1* mutants exhibited higher PSII effective quantum yields than wild type, attributable to reduced antenna size and enhanced carbon fixation capacity. The absence of NPQ accelerated high-light acclimation and was accompanied by increased xanthophyll accumulation, indicating that compensatory mechanisms can enhance overall photosynthetic efficiency. Together, these findings reveal the evolutionary origin of Lhcx/Lhcsr proteins and define the molecular basis of Lhcx1-mediated photoprotection in diatoms, providing fundamental insights into LHC-based photoprotection across photosynthetic lineages.

## Introduction

Photosynthesis consists of photochemical reactions in which two photosystems connected in series convert light energy into chemical energy. Among photosynthetic organisms, red-lineage secondary endosymbiotic algae, which possess plastids derived from red algae acquired through cellular endosymbiosis, are widely adapted to aquatic environments on Earth (Pierella Karlusich et al., 2020; Pierella Karlusich et al., 2025). Light conditions in these environments are as dynamic as those in terrestrial environments, making it extremely important for photosynthetic organisms to protect their photosystems from excess light to suppress the generation of harmful reactive oxygen species caused by the absorption of excessive light energy. Non-Photochemical Quenching (NPQ), which dissipates excess light energy as heat, plays a crucial role in this protective mechanism (Niyogi, 2000).

Multiple NPQ mechanisms with different induction times exist, and these exhibit distinct characteristics depending on the taxonomic group (Goss and Lepetit, 2015; Malnoë, 2018). In particular, energy-dependent quenching (qE) is a major component of NPQ and is carried out by Light-Harvesting Complex (LHC) proteins (Peers et al., 2009; Goss and Lepetit, 2015; Buck et al., 2019). In red-lineage secondary endosymbiotic algae including diatoms, qE depends on the Lhcx subfamily of the LHC family (Buck et al., 2019; Croteau et al., 2025). The diatom LHC is called FCP (Fucoxanthin-Chlorophyll *a*/*c*-binding Protein) because it binds pigments such as Chl *c*, fucoxanthin, diadinoxanthin, and diatoxanthin in addition to Chl *a*. In addition to Lhcx, xanthophyll cycle pigments play a crucial role in qE induction. In diatoms, the proton gradient (ΔpH) across the thylakoid membrane activates the diadinoxanthin to diatoxanthin converting cycle (Ddx-Dtx cycle) (Lavaud and Kroth, 2006; Goss and Lepetit, 2015). The violaxanthin-antheraxanthin-zeaxanthin (VAZ) cycle, well known in land plants (Niyogi et al., 1998), also functions in diatoms, complementing the Ddx-Dtx cycle (Lohr and Wilhelm, 1999; Giossi et al., 2025). Green algae, on the other hand, utilize qE dependent on the Lhcsr subfamily, which is homologous to Lhcx, as well as state transitions (qT), where LHCII detaches from photosystem II (PSII) and binds to photosystem I (PSI) through phosphorylation control to adjust the excitation balance between the two photosystems (Allorent et al., 2013; Minagawa and Tokutsu, 2015). Despite the well-established sequence homology between Lhcx and Lhcsr, these proteins exhibit differences in their regulatory mechanisms at the gene expression level (Allorent et al., 2016; Petroutsos et al., 2016; Tokutsu et al., 2019a; Tokutsu et al., 2019b; Im et al., 2024; Zhang et al., 2024). Furthermore, while land plants require PsbS for qE induction (Li et al., 2000; Niyogi, 2004), diatoms lack this protein. The evolutionary origin of Lhcx/Lhcsr has not been extensively discussed due to the lack of comprehensive phylogenetic analyses of LHC subfamilies.

NPQ in red-lineage algae has been studied in diatoms and *Nannochloropsis* belonging to heterokont algae (Bailleul et al., 2010; Buck et al., 2019; Park et al., 2019; Buck et al., 2021; Giovagnetti et al., 2022). In model diatom species, multiple Lhcx isoforms are expressed under different conditions and show functional redundancy. Therefore, almost complete Lhcx-deficient strains have not been created in diatoms to date, and the physiological responses and details of excitation energy transfer in states with significantly deficient NPQ have not been fully elucidated. In contrast, *Chaetoceros gracilis* has low redundancy of Lhcxs, with primarily only CgLhcx1 being expressed (Kumazawa et al., 2022), providing the possibility that an almost complete Lhcx-deficient strain can be obtained by creating a knockout of this CgLhcx1. Furthermore, the three-dimensional structures of PSII–FCPII (Nagao et al., 2019; Wang et al., 2019; Nagao et al., 2022) and PSI–FCPI (Nagao et al., 2020a; Xu et al., 2020) supercomplexes have already been elucidated in this species, establishing a foundation for structural biology research. This study aims to investigate the origin of Lhcx phylogenetically and to elucidate the mechanism of its excitation energy quenching for high light (HL) protection by creating Lhcx1-deficient mutants in *C. gracilis* using CRISPR/Cas9 genome editing technology.

## Results

### Origin of LHCs in the Lhcx/Lhcsr Subfamily

To elucidate the origins of the diatom Lhcx and green algal Lhcsr subfamilies, which perform NPQ, we conducted a comprehensive molecular phylogenetic analysis of light-harvesting complexes (LHCs) from red algae, green algae, cryptophytes, and diatoms, and represented the estimated phylogenetic tree as an unrooted tree (Figure 1). Comprehensive retrieval of LHC sequences by BLAST confirmed that, as in our previous reports, red algae possess only the Lhcr subfamily, cryptophytes have Lhcr and Lhcz subfamilies, and green algae contain Lhca, Lhcb, and Lhcsr subfamilies, while diatoms possess Lhcr, Lhcz, Lhcq, CgLhcr9 homolog, Lhcx, and Lhcf subfamilies (Kumazawa et al., 2022; Kumazawa and Ifuku, 2024). In Figure 1, each LHC subfamily in the red lineage is separated. However, within the Lhcr subfamily, there exists diversity among the Lhcr proteins from red algae, cryptophytes, and diatoms, encompassing gene groups with distant evolutionary relationships.

**Figure 1.**
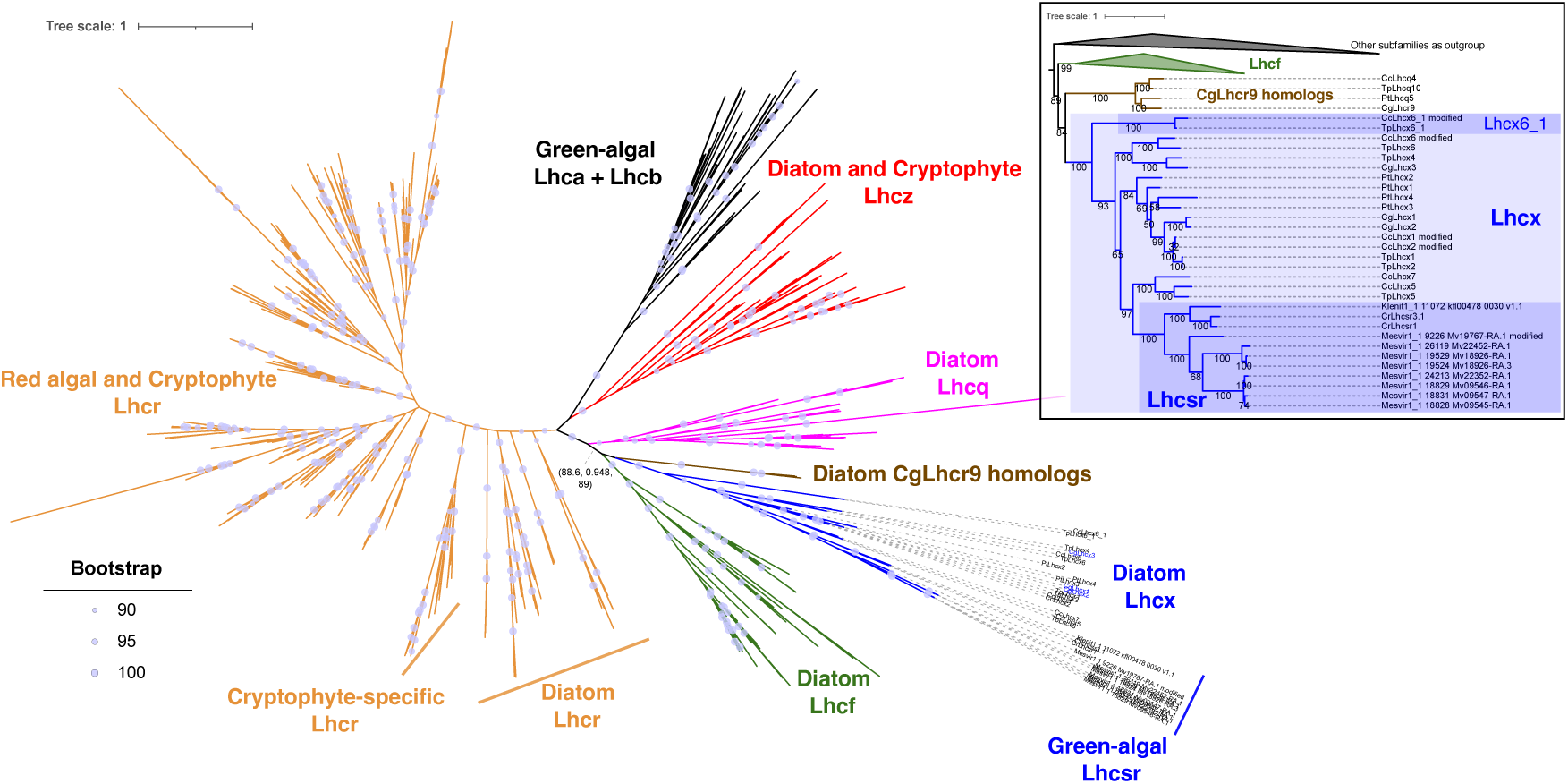
Molecular Phylogenetic Tree of Light-Harvesting Complexes (LHCs) from Red and Green Evolutionary Lineages. LHC sequences were obtained from red algae, green algae *Mesostigma viride* NIES-296 (Mesvir1_1) and *Klebsormidium nitens* NIES-2285 (Klepit1_1), diatoms: *Chaetoceros gracilis* (Cg), *Thalassiosira pseudonana* (Tp), *Cyclotella cryptica* (Cc), *Phaeodactylum tricornutum* (Pt), and cryptophytes: *Guillardia theta* CCMP2712 and Cryptophyceae sp. (*Baffinella* sp.) CCMP2293. These were combined with reference sequences for Lhcsr from *Chlamydomonas reinhardtii*: Lhcsr1 (UniProt: P93664) and Lhcsr3.1 (UniProt: P0DO19), which is identical to Lhcsr3.2 (UniProt: P0DO18), and a molecular phylogenetic tree was constructed using IQ-TREE2. The phylogenetic tree is displayed as an unrooted tree, and the inset tree is rerooted with the other LHC family members except for Lhcx, Lhcf, and CgLhcr9 homologs. LHC subfamilies, Lhcr, Lhcz, Lhcq, Lhcf, Lhcx, Lhca/Lhcb, and CgLhcr9 homologs, are indicated by orange, red, magenta, green, blue, black, and brown, respectively. Red algal and Cryptophyte Lhcrs are in the left part of the tree, Lhcrs expect for Cryptophyte-specific Lhcrs and Diatom Lhcrs. Detailed annotations are shown only for the Lhcx/Lhcsr subfamily. Circles on nodes indicate ultra-fast bootstrap support values of 90% or higher. Numbers in parentheses indicate SH-aLRT support values (%) / aBayes support values / ultra-fast bootstrap support values (%). Numbers in the inset tree are ultra-fast bootstrap support values (%).

Lhcx and Lhcsr formed a monophyletic clade with an ultrafast bootstrap value of 100% (Figure 1, inset). Within this clade, *Thalassiosira pseudonana* Lhcx6_1 and *Cyclotella cryptica* Lhcx6_1, derived from diatoms belonging to the order Thalassiosirales, clearly branched at the most basal position and were evolutionarily distant from typical Lhcx and Lhcsr proteins. All Lhcx1–3 proteins from the diatom *Chaetoceros gracilis* were positioned within the typical Lhcx clade. Additionally, the Lhcx and Lhcsr clade showed monophyly with the Lhcq, CgLhcr9 homolog, and Lhcf subfamilies, supported by a high bootstrap value (100%). Within that clade, Lhcq was located at the most basal position, while the Lhcx/Lhcsr clade was positioned internally. In contrast, the Lhcx/Lhcsr clade did not show monophyly with the Lhca/Lhcb clade. Furthermore, the Lhcr of red algae and the Lhca and Lhcb subfamilies, which are responsible for light harvesting in red and green algae possessing primary plastids, did not form a monophyletic group; between them existed cryptophyte Lhcr, diatom Lhcr, cryptophyte and diatom Lhcz, as well as other subfamilies of the red evolutionary lineage. These data suggest that Lhcsr and Lhcx subfamilies are derived from the red-lineage specific subfamilies.

### Creating *Lhcx1* Knockout Strains with Ribozyme-Based Genome Editing

*C. gracilis* has three typical Lhcx genes (*CgLhcx1–3*): according to the genome database of *C. gracilis* (ChaetoBase v1.1, https://chaetoceros.nibb.ac.jp/), *CgLhcx1* showed the highest transcription level by a significant margin (Kumazawa et al., 2022). Therefore, we decided to create an *Lhcx1*-deficient mutant in *C. gracilis* using the CRISPR/Cas9 system. To facilitate the comprehensive analysis of LHC genes using genome-editing in *C. gracilis,* we have developed the inducible CRISPR/Cas9 system, in which both Cas9 and guide RNA (gRNA) expression is regulated by the inducible Pol-II-type promoter. First, we introduced the *Cas9* gene under the nitrate reductase (NR) promoter into wild-type *C. gracilis* with a vector having the nourseothricin resistance gene (*nat*) via electroporation (Supplemental Figure 1A, B). Colonies obtained through antibiotic-resistance selection were further selected for high Cas9 expression in the presence of nitrate by immunoblotting (Supplemental Figure 1B). The selected hC9_5 strain showed morphology and growth characteristics equivalent to the wild-type strain (WT) and was established as the host strain for the genome editing (Supplemental Figure 1C, D). Second, to express mature gRNA in the host hC9_5 under the NR promoter, we employed a method utilizing ribozymes (Gao and Zhao, 2014; Zhang et al., 2017; Taparia et al., 2022). This method employed a Ribozyme-gRNA-Ribozyme (RGR) construct in which a hammerhead ribozyme with self-cleavage activity at the 3’ end was attached to the 5’ end of the gRNA, and an HDV ribozyme with self-cleavage activity at the 5’ end was attached to the 3’ end of the gRNA. This RGR cassette for the gene of interest can be easily constructed by overlap PCR and inserted into pCgNRp^ble^ vector, having an antibiotic-resistant gene against zeocin (*ble*) for the second selection.

The transformation of the host hC9_5 strain with the linearized pCgNRp^ble^-Lhcx1-RGR vector produced 252 zeocin-resistant colonies, from which 23 strains were randomly selected and 13 strains had various mutations in the target sequence of the *CgLhcx1* gene (Supplemental Figure 2A, B, C). Among the strains confirmed for target sequence editing, *lhcx1-7*, *15*, and *234* strains were used for subsequent analysis. When the nucleotide sequences of the *Lhcx1* region were confirmed by Sanger sequencing, heterogeneity was observed (Supplemental Figure 2C). Since *C. gracilis* is a diploid organism like other diatoms, mutagenic events can occur in both alleles to completely inactivate the target gene. However, simultaneous mutations in both alleles are rare, and in most cases only one allele is mutated or neutral mutations such as in-frame deletions/insertions may occur, resulting in the formation of mixed cell populations with different mutation status and types through subsequent cell divisions (Huang and Daboussi, 2017). Therefore, we examined the accumulation of Lhcx1 protein in these strains by western blotting using an anti-Lhcx1 antibody (Supplemental Figure 2D). Almost no accumulation of Lhcx1 protein was observed in the *lhcx1-7* and *lhcx1-15* strains. On the other hand, the *lhcx1-234* strain showed reduced accumulation of Lhcx1 compared to WT, but it was not a complete deficiency since *lhcx1-234* may include neutral mutations such as in-frame deletions/insertions.

### Phenotypic Analysis of *Lhcx1* Deficient Strains

WT and *lhcx1-7*, *15*, and *234* strains were cultured with air bubbling for 7–8 days under low light (LL) at 30 μmol photons m^−2^ s^−1^ and high light (HL) at 300 μmol photons m^−2^ s^−1^. We selected 300 μmol photons m^−2^ s^−1^ as high light because it is still much higher than the light intensity at the deep chlorophyll maximum layer, where diatoms are typically most abundant. No significant differences in growth were observed between WT and genome-edited strains (Supplemental Figure 3). To investigate the photosynthetic characteristics of WT and *lhcx1* strains, pulse-amplified modulated (PAM) fluorescence measurements were performed on day 3 and day 7–8 using cells cultured under LL and HL conditions (Figure 2, Supplemental Figure 4). As a result, *lhcx1-7*, *15*, and *234* were hardly able to induce NPQ, an indicator of thermal dissipation of excess excitation energy around PSII, under both LL and HL conditions (Figure 2A, Supplemental Figure 4A). The *lhcx1* strains cultured under LL showed lower photochemical quenching qP under light exposure compared to WT, indicating a more reduced state of the plastquinone pool. However, this trend was not clear in the *lhcx1* strains cultured under HL, because non-regulated energy dissipation due to photoinhibition affects the qP level (Figure 2B). The effective quantum yield of PSII, Y(II), showed higher values in all *lhcx1* strains compared to the WT strain in cells cultured under HL for 7–8 days (Figure 2C). In cells cultured under HL for 3 days, *lhcx1* strains showed a trend of higher Y(II) compared to WT, but the difference was not significant (Supplemental Figure 4C). The *lhcx1-7, 15* strains exhibited significantly higher oxygen-evolving activity compared to WT cells at the same total Chl *a* content of 5 μg (Figure 3). The *lhcx1-234* strains showed the tendency towards higher oxygen-evolving activity, though it was not statistically significant (*p* = 0.157). This may be due to the mosaic genetic background of *Lhcx1* mutations in *lhcx1-234*.

**Figure 2.**
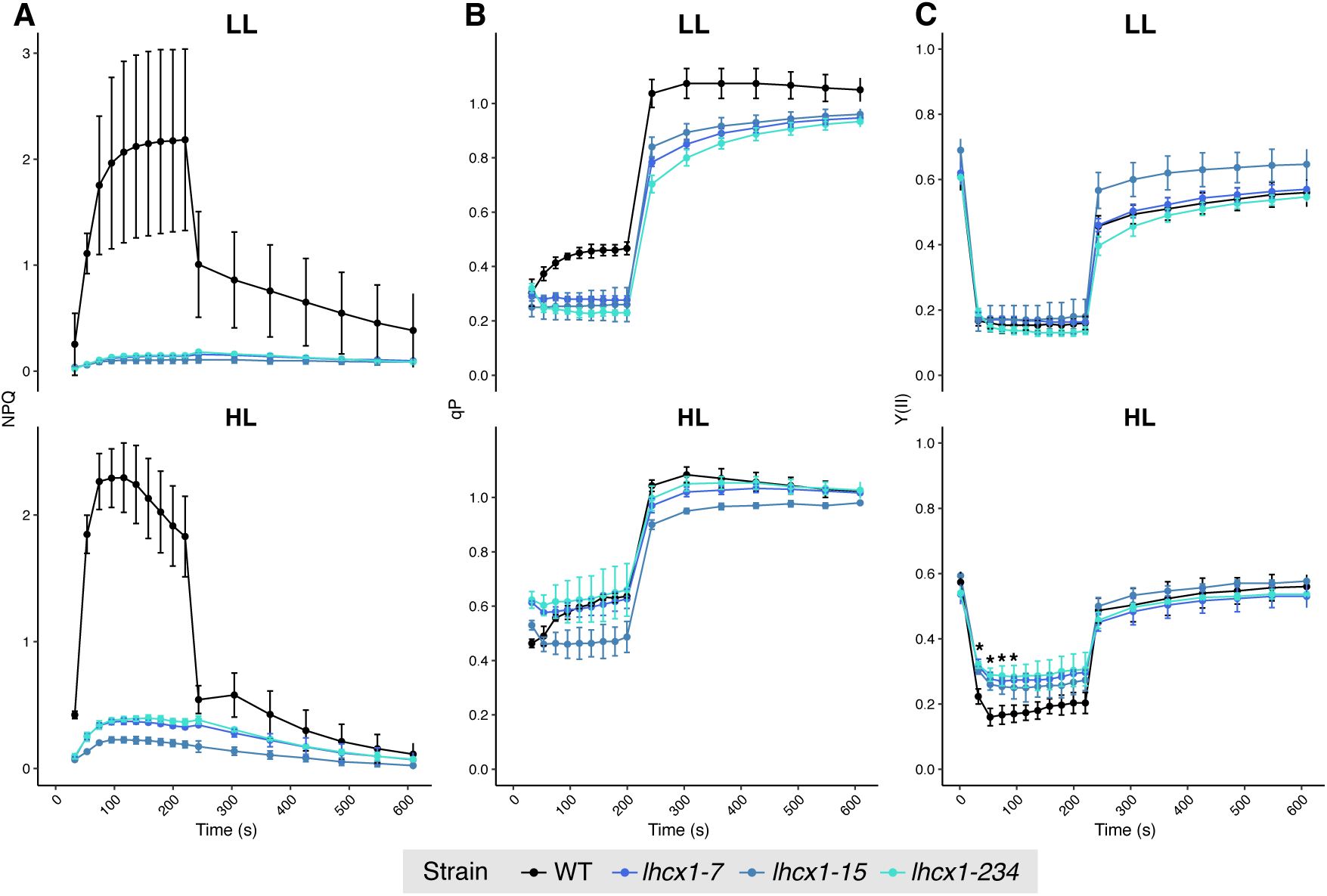
PAM Fluorescence Measurements of *C. gracilis* Wild-Type and *lhcx1* Strains Cultured with Aeration for 7–8 Days Under Low Light (LL) and High Light (HL) A) NPQ of LL and HL cultures. B) qP of LL and HL cultures. C) PSII effective quantum yield Y(II) of LL and HL cultures. * indicates that adjusted *p* < 0.05 by Tukey’s HSD test was satisfied between the wild-type and all *lhcx1* strains. The points in each figure represent mean ± standard deviation with biological replicates of *n* = 3.

**Figure 3.**
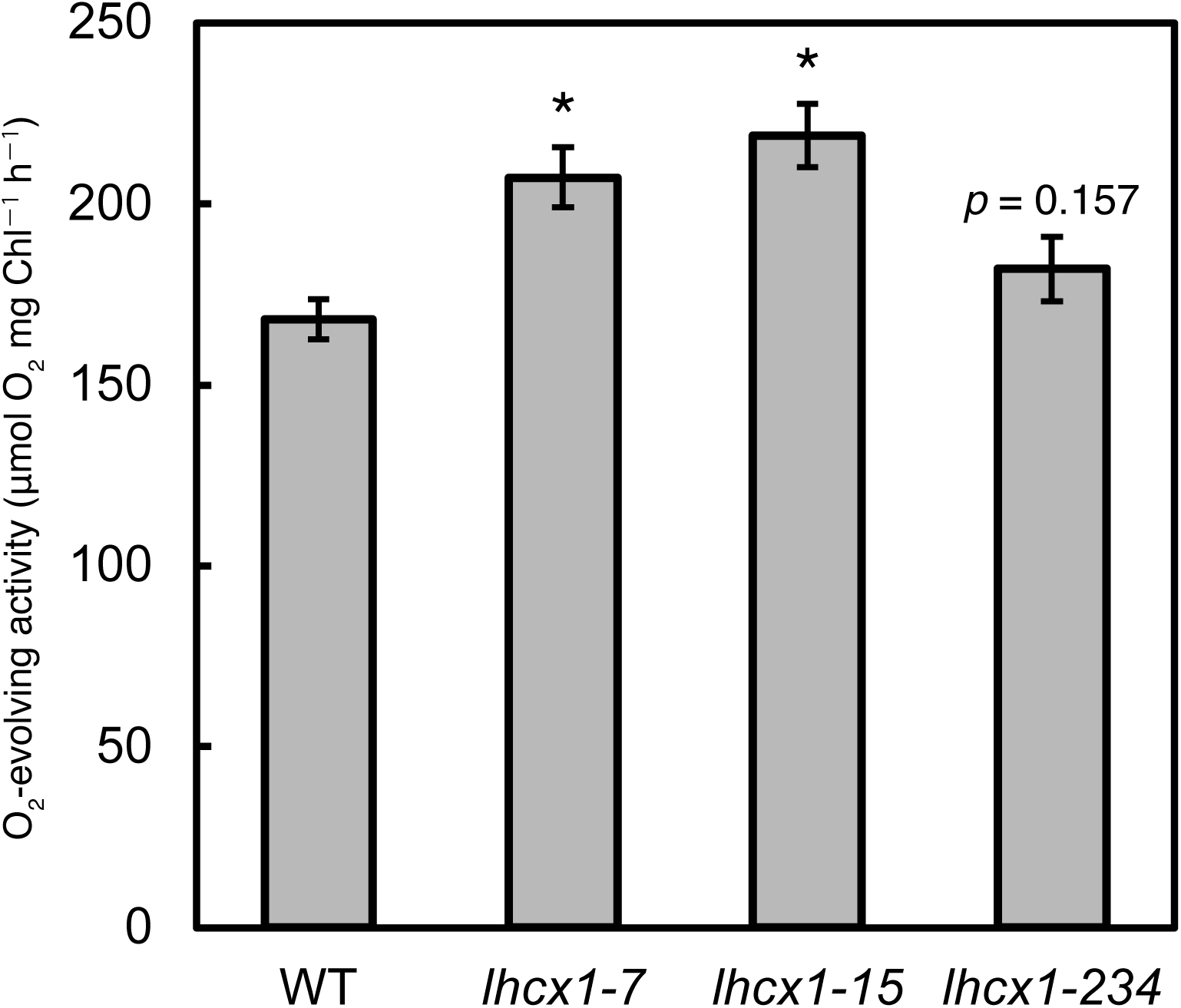
Oxygen-Evolving Activity of *C. gracilis* Wild-Type and *lhcx1* Strains. This figure shows the oxygen-evolving activity of *C. gracilis* wild-type (WT) and three *lhcx1* mutant strains (*lhcx1-7*, *lhcx1-15*, and *lhcx1-234*). Measurements were conducted under a light intensity of 300 μmol photons m^−2^ s^−1^ in the presence of 10 mM NaHCO_3_. Data are presented as mean ± standard deviation (*n* = 3). Statistical analysis was performed using Dunnett’s test to compare each mutant strain with the wild-type (WT). Asterisks (*) indicate p<0.001, while other actual p-values are shown in the graph.

### Whole-Cell Absorption Spectra and Pigment Analysis

The whole-cell absorption spectra showed no significant differences between WT and *lhcx1* mutants in cells cultured under LL, while in the cells cultured under HL, the *lhcx1* mutants exhibited greater absorption in the 400–500 nm region compared to the WT (Supplemental Figure 5). Since this region corresponds to carotenoid absorption, pigment analysis using HPLC (High Performance Liquid Chromatography) was performed to examine carotenoid contents. The amount of Chl *a* per cell was decreased in the mutants compared to WT under LL conditions (Figure 4A). Additionally, under HL conditions, WT also showed a decrease in Chl *a* content compared to LL conditions, but the decrease tends to be greater in the *lhcx1* mutants. Since the total absorption spectra were normalized at the Qy peak of Chl *a*, carotenoid content was quantified per Chl *a* amount. Fucoxanthin, the major carotenoid in *C. gracilis* FCP, showed no significant differences between WT and mutants under both light conditions, except for *lhcx1-15* under HL conditions (Figure 4B). The *lhcx1-15* under HL conditions showed slightly lower fucoxanthin content compared to WT under the same conditions. On the other hand, for diadinoxanthin (Ddx) and diatoxanthin (Dtx), the major pigments of the diatom xanthophyll cycle, no significant differences in accumulation were observed between WT and *lhcx1* mutants under LL conditions (Figure 4C, D). In contrast, under HL conditions, WT showed greater accumulation of Dtx compared to LL conditions. The mutant strains accumulated significantly more Dtx compared to the WT under the same conditions. Regarding Ddx, both WT and mutants showed greater accumulation under HL conditions compared to LL conditions, but significant increases in accumulation in mutants compared to WT were not observed, except for *lhcx1-15*. Total xanthophyll cycle pigments (Ddx+Dtx) were also increased in *lhcx1* mutants under HL (Figure 4E). The de-epoxidation state (DEPS = [Dtx]/([Dtx]+[Ddx])), an indicator of the xanthophyll cycle, was calculated from the accumulation of Dtx and Ddx (Figure 4F). Under LL, there was no significant difference between WT and mutants. Under HL, the WT showed a higher DEPS (0.34) compared to LL conditions. The mutants showed significantly higher DEPS (0.47–0.52) compared to WT under the same conditions.

**Figure 4.**
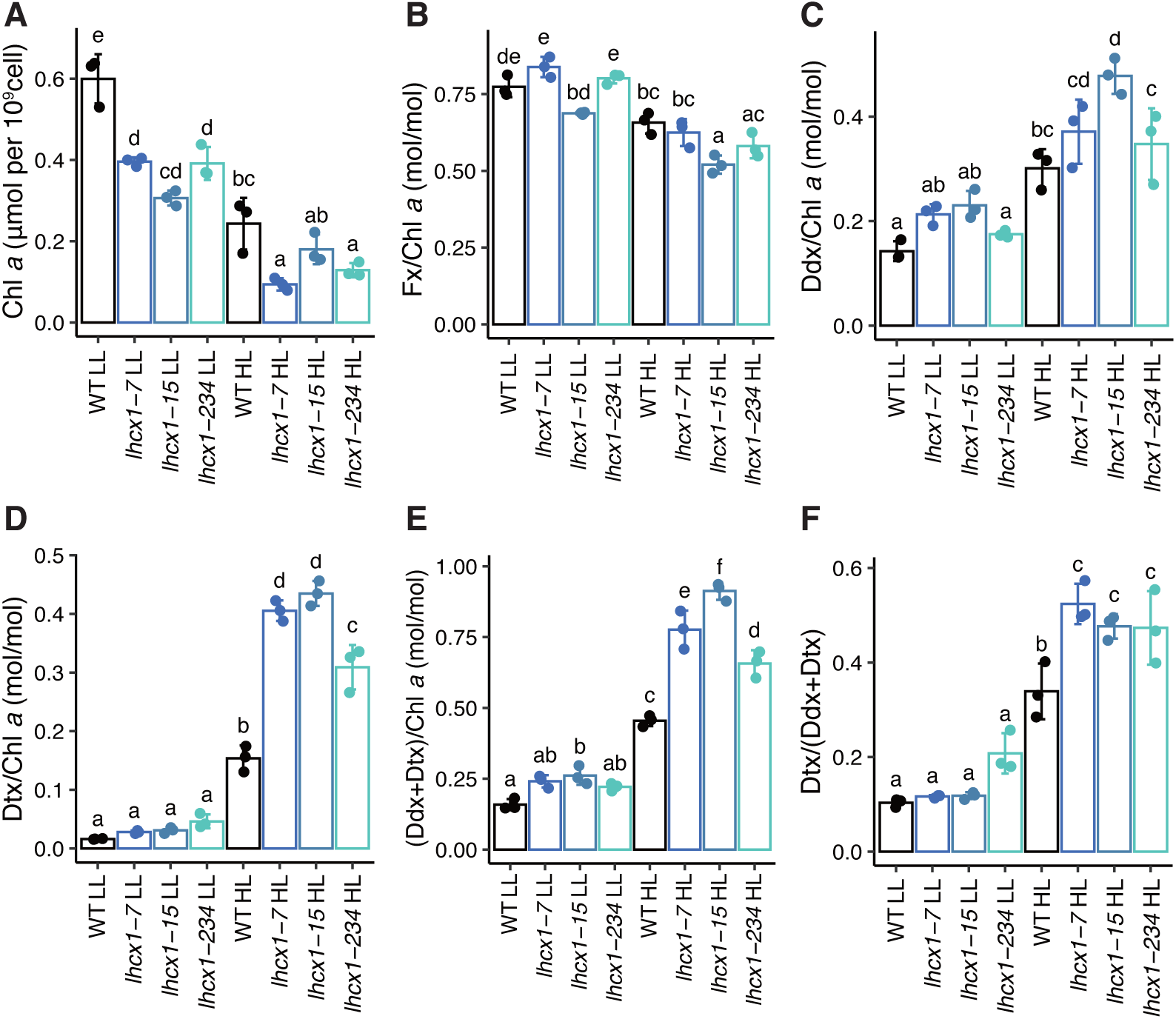
Pigment Analysis of Low Light (LL) and High Light (HL) Cultures. A) Chlorophyll (Chl) *a* content per 10^9^ cells. B) Fucoxanthin (Fx) content per Diadinoxanthin (Ddx) content per Chl *a*. D) Diatoxanthin (Dtx) content per Chl *a*. E) Ddx+Dtx content per Chl *a*. F) De-epoxidation state (DEPS = [Dtx]/([Dtx]+[Ddx])). Data are presented as mean ± standard deviation (*n* = 3). Different alphabets indicate adjusted *p* < 0.05 by Tukey’s HSD test.

### Steady-State Low-Temperature Fluorescence and Antenna Size Measurements

WT and *lhcx1* cultured under HL were subjected to 30-minute treatments under either dark conditions (dark-treated) or HL conditions (HL-treated), then were immediately frozen in liquid nitrogen, and used for the low-temperature (77K) fluorescence spectra measurements (Figure 5A, B). WT was measured as two independent samples cultured separately (solid and dashed lines in Figure 5A, B). The 77K steady-state fluorescence spectra well represent the distribution of excitation energy between photosystems and light-harvesting pigment proteins. The excitation wavelength was 459 nm, which efficiently excites Chl *c* and fucoxanthin bound to the peripheral antenna FCP in diatoms. The 77K fluorescence spectra of *C. gracilis* showed two peaks derived from CP43 and CP47 of photosystem II, observed around 686 nm and 694 nm, respectively (Andrizhiyevskaya et al., 2005; Nagao et al., 2010). In all cells cultured under HL, the mutants showed lower CP43 peaks compared to the WT in spectra normalized at the CP47 peak (Figure 5A). The fluorescence intensity around 680 nm, which is considered to be the fluorescence from FCP, was smaller in *lhcx1* mutants under dark conditions compared to the WT. These indicate that energy transfer from the peripheral antenna FCP via CP43 is prominent in the dark especially in WT. This also suggests that the effective antenna size of PSII differed between WT and *lhcx1* in the dark. Therefore, the antenna size after dark adaptation of cells cultured under HL was measured by measuring the induction curve of Chl fluorescence with a Fluorometer FL 3500 (PSI, Czech Republic) (Nedbal et al., 1999). As a result, it was revealed that *lhcx1* had a significantly smaller effective antenna size of PSII compared to WT in all strains (Supplemental Figure 6).

**Figure 5.**
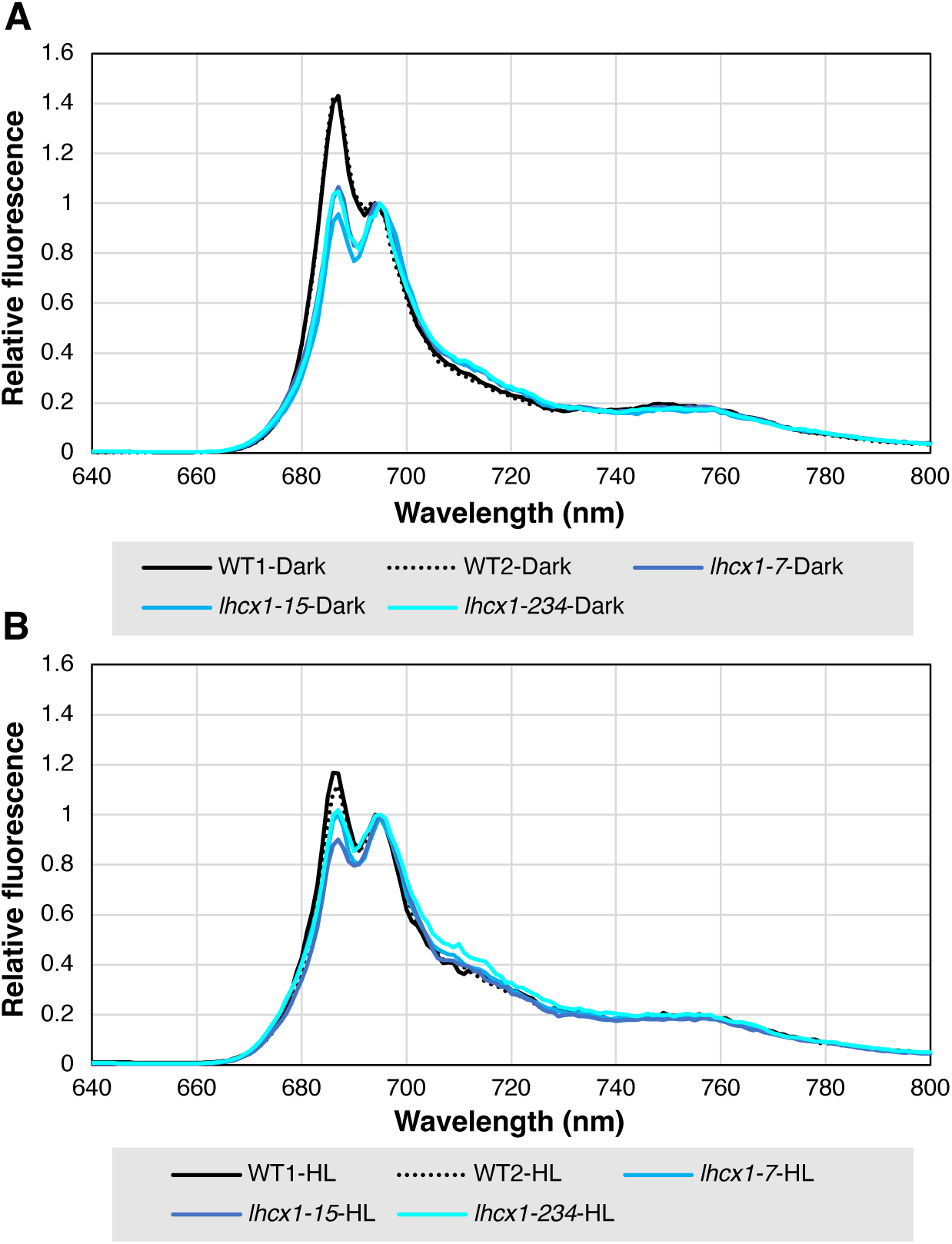
Low-Temperature Fluorescence Emission Spectra of Wild-Type (WT) and *lhcx1* Mutants Cultured Under High Light. A) Low-temperature fluorescence emission spectra measured after 30 minutes of dark treatment followed by rapid freezing in liquid nitrogen. B) Low-temperature fluorescence emission spectra measured after 30 minutes of HL treatment followed by rapid freezing in liquid nitrogen. Each spectrum was normalized at the peak corresponding to CP47 (693–695 nm). The wild type was measured in two separately cultured samples (black solid line and black dashed line).

### Time-Resolved Low-Temperature Fluorescence Spectra

To elucidate the process of excitation energy transfer from the peripheral antenna FCP to the photosystems, time-resolved fluorescence (TRF) spectra were measured under the same conditions as the samples subjected to low-temperature steady-state fluorescence spectrum measurements. TRF fluorescence was measured on samples of whole cells that were frozen in liquid nitrogen immediately after dark treatment and HL treatment (300 μmol photons m^−2^ s^−1^). This allows observation of excitation energy transfer and quenching in states where NPQ is not induced and where it is induced (Chukhutsina et al., 2017). Fluorescence decay-associated spectra (FDAS) were obtained by global analysis that decomposed the obtained TRF spectra into multiple exponential components with time as a variable (Figure 6, Supplemental Figure 7). In FDAS, positive peaks (decay) indicate the decay of fluorescence, while negative peaks (rise) indicate the rise of fluorescence. From the TRF spectra of the WT, FDAS were constructed with six-time constants. The assignment of each peak followed previous literature (Andrizhiyevskaya et al., 2005; Nagao et al., 2010). In the dark-adapted WT, the FDAS with a lifetime of 40 ps showed positive peaks of FCP at <680 nm and a large negative peak around 687 nm corresponding to CP43 (black solid line in Figure 6 and Supplemental Figure 7). This reflects excitation energy transfer from FCP to CP43. In the spectrum of the second lifetime of 140 ps, positive peaks were observed not only at ∼680 nm but also around 687 nm corresponding to CP43, while the waveform around 694 nm corresponding to CP47 showed a convex downward shape. This is thought to be due to the negative peak of CP47 overlapping with the shoulder of the nearby positive peak of CP43, and is considered to indicate excitation energy transfer from FCP/CP43 to CP47. Also, in the second lifetime component, a PSI positive peak at ∼710 nm was observed, which, together with the negative peak of FCP, indicates the excitation energy transfer from FCP to PSI. The third lifetime of 640 ps continued to show the same trend as 140 ps, suggesting that excitation energy transfer from FCP/CP43 to CP47 was ongoing. The fourth and fifth lifetimes, at 1.6 ns and 3.8 ns, showed positive peaks corresponding to CP43 and CP47, indicating simple fluorescence decay. The longest lifetime of 26 ns did not exhibit a prominent positive peak around 710 nm, which is characteristic of PSI fluorescence, but instead showed positive peaks corresponding to CP43 and CP47, associated with charge recombination in photosystem II. Therefore, excitation energy transfer from PSII to PSI, so-called spillover, did not occur.

**Figure 6.**
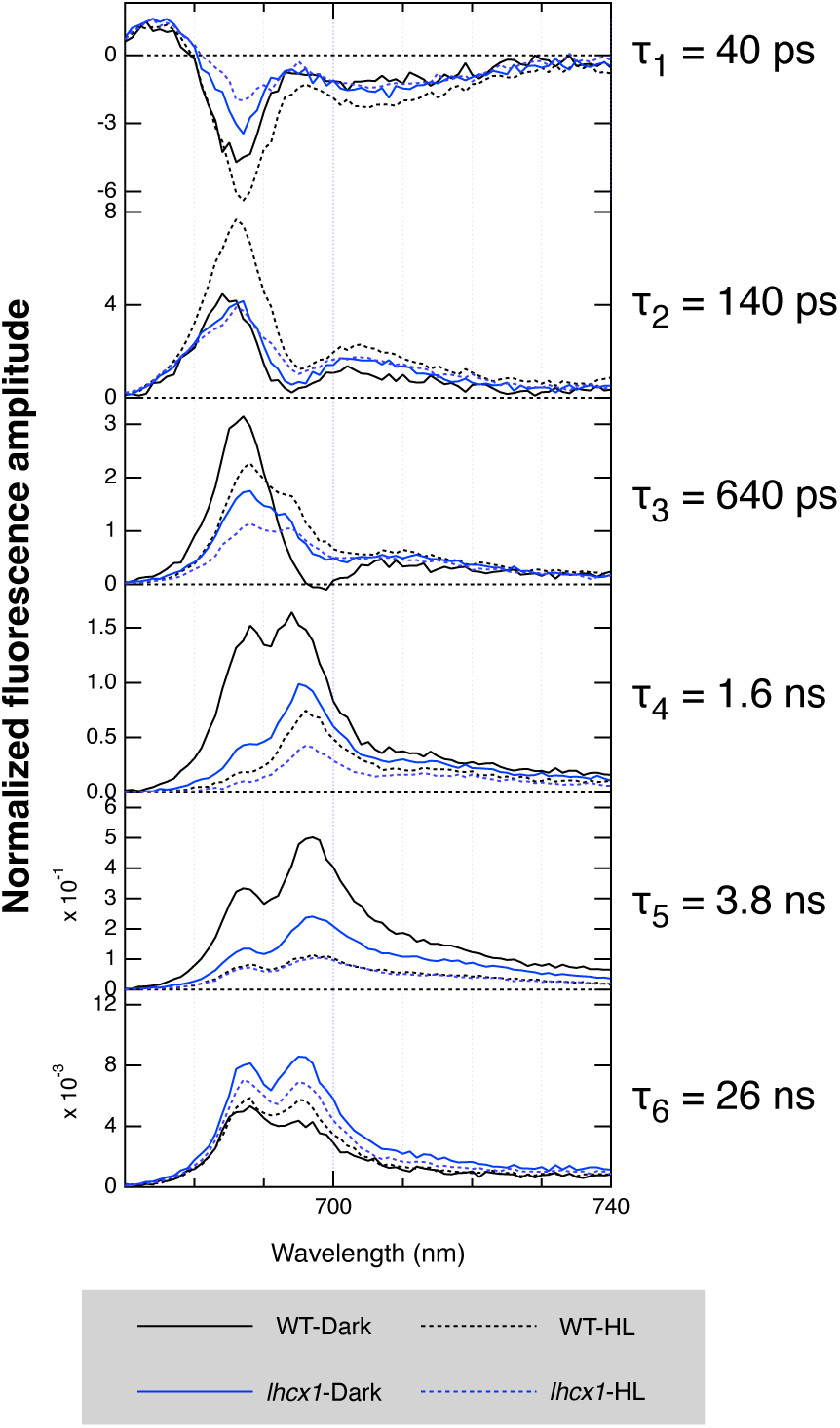
Fluorescence Decay-Associated Spectra of Time-Resolved Fluorescence. Black lines indicate WT, blue lines indicate *lhcx1-7*, solid lines represent measurements after 30 minutes of dark treatment, and dashed lines represent measurements after 30 minutes of treatment at 300 μmol photons m^−2^ s^−1^.

On the other hand, WT subjected to HL (black dashed line in Figure 6 and Supplemental Figure 7) showed a similar waveform at 40 ps as the dark-adapted WT. At 140 ps, although the positive peak around 687 nm was much larger compared to the dark-adapted WT, the value at the dip around 694 nm was larger than that of the dark-adapted WT. In other words, WT acclimated to HL had a smaller fluorescence rise at CP47 compared to the dark-adapted sample, while the energy decay around 687 nm was larger. The *lhcx1* mutant, which does not induce NPQ, showed spectra similar to the dark-adapted WT in the first- and second-time constants when dark-adapted (blue, green, and red solid lines in Figure 6 and Supplemental Figure 7A, B). However, in the third time constant (640 ps), while the dark-adapted WT showed excitation energy transfer from FCP/CP43 to CP47, *lhcx1* already showed positive peaks at both 687 nm and 694 nm, indicating fluorescence decay of CP43 and CP47. In subsequent time constants, 687 nm and 694 nm also showed only positive peaks. The HL-acclimated *lhcx1* (blue, green and red dashed line in Figure 6 and Supplemental Figure 7A, B) also showed spectra similar to the dark-adapted *lhcx1* at all time constants and did not show the large positive peak around 687 nm at 140 ps observed in the HL-acclimated WT. Furthermore, since energy transfer from FCPII to CP43 and CP47, and from CP43 to CP47 was completed by 140 ps, similar to the HL-treated WT and dark-treated *lhcx1*, it was concluded that *lhcx1* has a smaller antenna size compared to the WT even under HL conditions. From the above, while the decay of fluorescence around 687 nm is normally attributed to energy transfer from CP43, when looking at the energy transfer of *lhcx1* at the same time constant, such decay is not observed. Therefore, it is concluded that the decay of fluorescence around 687 nm in the HL-acclimated WT is not decay of CP43 but quenching attributed to Lhcx1.

### Molecular interaction of Lhcx1 in thylakoid membranes

To determine the localization of Lhcx, we performed native-PAGE followed by second-dimension SDS-PAGE and subsequent immunoblotting using anti-Lhcx1 antibody. Since conventional Clear Native (CN)-PAGE with deoxycholate made it difficult to separate diatom C_2_S_2_M_2_ (Core_2_–S-tetramer_2_–M-tetramer_2_) PSII–FCPII while keeping FCP-A (which contains CgLhcf1 as a major component) bound to PSII (Nagao et al., 2012), we applied a modified CN-PAGE using Amphipol A8-35 (Kameo et al., 2021), which enhances the stability of solubilized protein complexes, instead of deoxycholate as the surfactant that imparts negative charge to proteins (Figure 7A). In the diatom *C. gracilis*, a dense band was observed at a higher molecular weight position than spinach C_2_S_2_M_2_ (Core_2_–S-trimer_2_–M-trimer_2_) PSII–LHCII. Additionally, a broad brown band, which was observed at a higher molecular weight position than the LHC/FCP monomer, extended to a lower molecular weight position than the spinach LHCII trimer. This CN-PAGE gel strip was subjected to SDS-PAGE with a gel containing 6 mol L^−1^ urea to separate each subunit (Figure 7B). Annotation of each spot was based on previous reports (Nagao et al., 2007; Zhou et al., 2024). The amino-acid sequence of HGRIAQLAFLGN shared with CgLhcf2 and CgLhcf12 was identified from the major FCP fraction (Ishihara et al., 2015). These reports indicate that FCP-B/C is one of the major FCPs in *C. gracilis*. Furthermore, since the FCP-B/C heterodimer is presumed to be loosely bound to C_2_S_2_M_2_ PSII–FCPII and transfer excitation energy, it was designated as L (Loose)-dimer (Figure 7A, B). The thylakoid membranes of *lhcx1* were also subjected to CN-PAGE (Supplemental Figure YYY). While *lhcx1* has the C_2_S_2_M_2_ PSII–FCPII as well as WT, *lhcx1* has a slightly paler broad band of L-dimers. Additionally, immunoblotting was performed to identify the Lhcx1 spot from the spots in the two-dimensional SDS-PAGE (Figure 7C). The Lhcx1 spot, indicated by the blue arrowhead in Figure 7C, was detected at a higher molecular weight than FCP-B/C in 2D-SDS-PAGE. In the 1D-CN-PAGE direction, the spot of Lhcx1 was detected across a wide range of multimers from monomers to higher-order multimeric complexes, co-migrating with FCP-B/C (Figure 7B). That spot of Lhcx1 in 2D-PAGE with oriole stain is paler than FCP-A/B/C because of lower accumulation of Lhcx1 than FCP-A/B/C. This result suggests that Lhcx1 may interact with the FCP-B/C dimers, which function as the loosely associated antenna of C_2_S_2_M_2_ PSII–FCPII.

**Figure 7.**
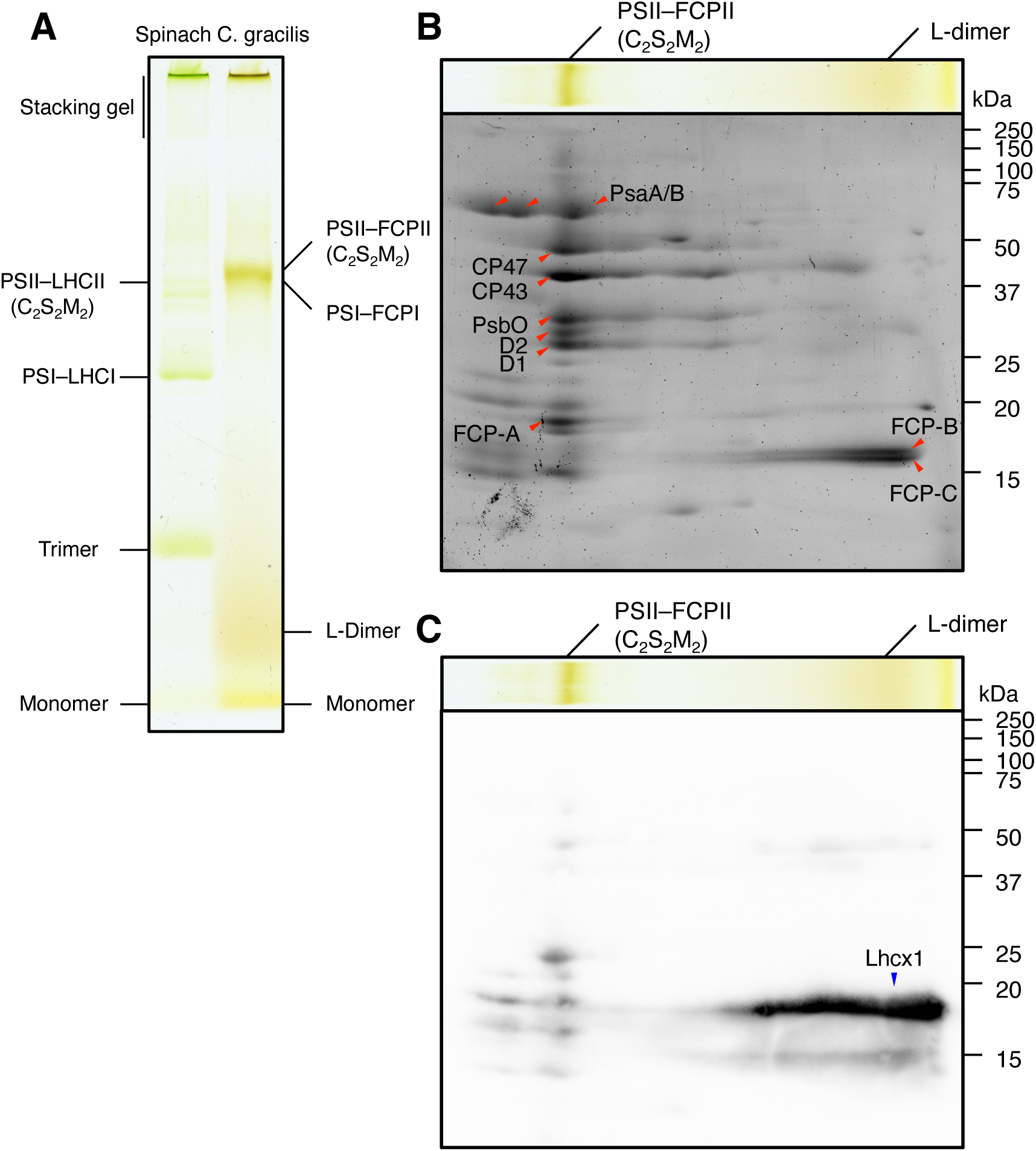
2D-Amphipol-based-CN/SDS-PAGE of *C. gracilis* Thylakoid Membranes. A) Amphipol-based CN-PAGE of *C. gracilis* and spinach thylakoid membranes. Annotation of each band for spinach follows previous reports (Järvi et al., 2011; Kameo et al., 2021). B) Protein fluorescence staining image of two-dimensional electrophoresis of Amphipol-based CN-PAGE. Annotation of each spot indicated by red arrowheads is based on previous reports (Nagao et al., 2007). FCP-A, FCP-B, and FCP-C correspond to CgLhcf1, CgLhcf12, and CgLhcf2, respectively (Nagao et al., 2019; Nagao et al., 2022; Zhou et al., 2024). C) Identification of the Lhcx1 spot by immunoblotting using anti-Lhcx1 antibody. The blue arrowhead indicates the detected spot.

### Phenotype of *lhcx1* Mutants in High CO_2_ Environment

We investigated whether the *lhcx1* mutants, which showed a higher effective quantum yield of PSII when acclimated to HL under normal atmospheric aeration conditions, would further improve their PSII effective quantum yield by enhancing downstream electron sink capacity with 3% CO_2_ addition. First, even when cultured with CO_2_ aeration, *lhcx1* showed almost no NPQ regardless of light intensity (Supplemental Figure 9A). The effective quantum yield of PSII, Y(II), increased in *lhcx1* when cultured with CO_2_ aeration even under LL conditions, where no difference was observed with atmospheric aeration (solid line in Supplemental Figure 9C). This Y(II) under light irradiation increased with time during the initial irradiation period and reached a constant value after about 2 minutes, which is thought to be due to the activation of the carbon assimilation pathway. Additionally, an increase in Y(II) was also observed under HL conditions. Photochemical quenching qP, which indicates the redox state of plastoquinone, was similar between WT and *lhcx1* under LL conditions. As observed in Figure 2B, qP of *lhcx1* varied among strains, with strains having lower Y(II) showing lower qP under HL conditions (Supplemental Figure 9B).

To understand the general physiological response of *lhcx1*, we performed RNA sequencing analysis under LL/HL and ambient/3%-CO_2_-supplemented air conditions. Under ambient air conditions, HL treatment induced 1,494 genes in WT and 1,530 genes in *lhcx1* mutant, with 458 genes commonly upregulated in both genotypes. Under CO_2_-enriched conditions, HL induced substantially more genes: 2,411 in WT and 2,402 in *lhcx1* mutant. To identify responses compensating the absence of NPQ in *lhcx1*, we compared the magnitude of HL response (fold-change from LL to HL) between genotypes. *lhcx1* showed enhanced HL response compared to WT in 978 genes under ambient air and 311 genes under CO_2_-enriched conditions, with 105 genes showing this enhanced response under both ambient and CO_2_ conditions. KEGG pathway enrichment analysis reveals that while both WT and *lhcx1* respond to HL through porphyrin metabolism and photosynthesis pathways (Figure 8A), *lhcx1* mutant specifically enhances carbon fixation and central carbon metabolism pathways under both ambient air and CO_2_ conditions (Figure 8B).

**Figure 8.**
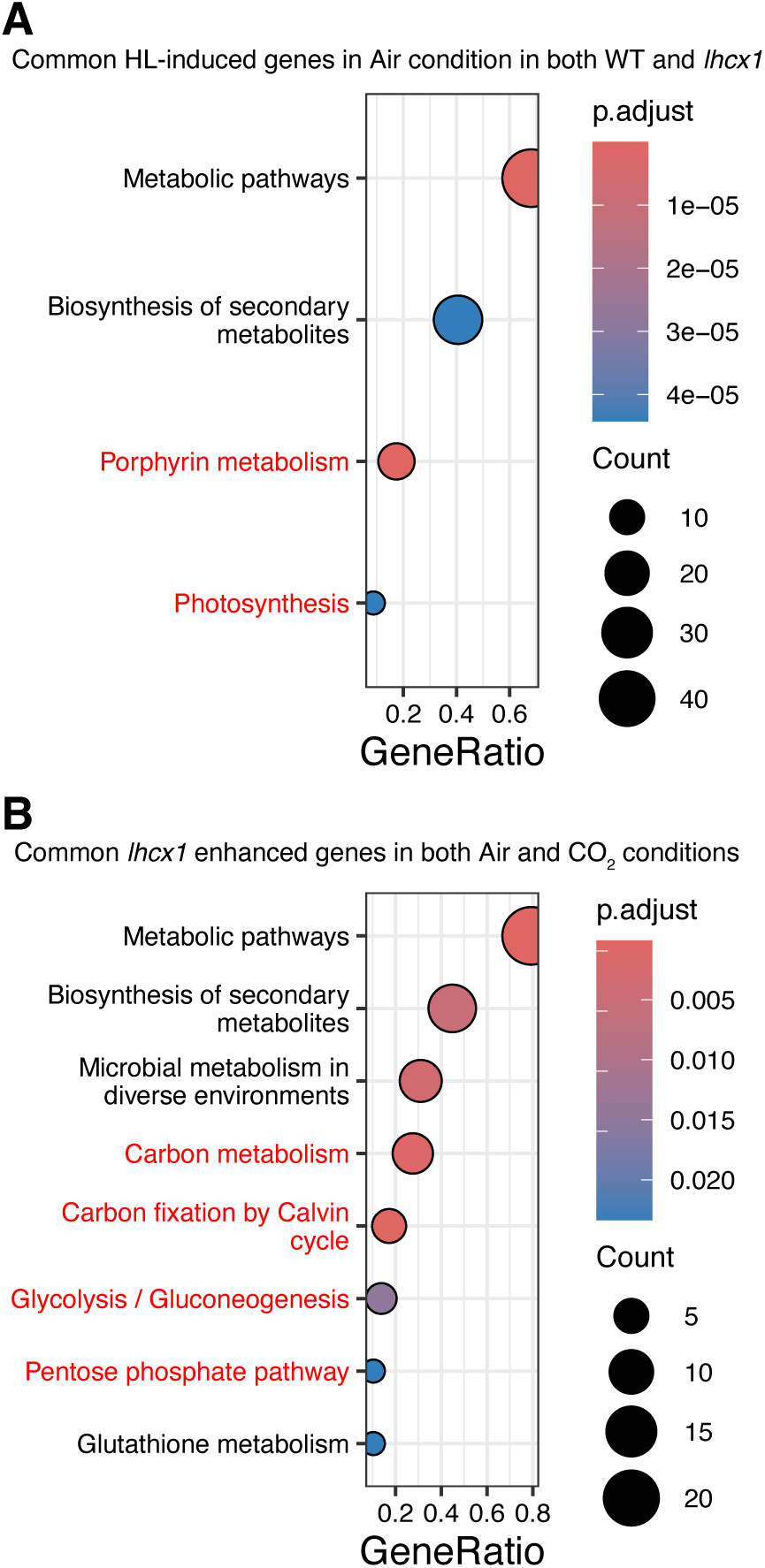
Gene Expression Analysis under CO_2_ Enrichment. KEGG pathway enrichment analysis of differentially expressed genes in the *lhcx1* mutant under LL and HL conditions. A) Pathways enriched in genes commonly induced by HL in both WT and the *lhcx1* mutant under ambient air conditions. B) Pathways enriched in genes showing enhanced HL response in *lhcx1* mutant under both ambient air and CO_2_-enriched conditions. Dot size represents gene count, and color intensity indicates statistical significance (q-value shown as p.adjust). Pathways related to typical HL response and photosynthesis are indicated with red letters.

## Discussions

The deficiency of Lhcx, while Lhcx belongs to the LHC family primarily responsible for light harvesting, almost completely suppresses the induction of non-photochemical quenching (NPQ). The NPQ-deficient *lhcx1* strains acclimated by inducing various HL defense systems possessed by *C. gracilis* and showed higher PSII activity compared to the WT. Since the increase in PSII activity was also induced in *lhcx1* with the addition of CO_2_, it was revealed that acclimation of downstream electron sinks in the electron transport chain, along with acclimation of the upstream light-harvesting system, plays an important role in HL acclimation of *C. gracilis*. This study suggests that the multiple photoprotection mechanisms can be interrelated and flexibly regulated in response to environmental changes in the marine diatom *C. gracilis*.

### Origin of Lhcx

The Lhcx subfamily, which is responsible for non-photochemical quenching (NPQ) in red-lineage secondary endosymbiotic algae such as heterokonts including diatoms, and presumably haptophytes, and the Lhcsr subfamily, which similarly mediates NPQ in green algae, streptophyte algae, and bryophytes, have been considered to share a common ancestral gene as they form a monophyletic group (Koziol et al., 2007). Although the Lhcx/Lhcsr subfamily is suggested to be derived from the red lineage, detailed molecular phylogenetic relationships with other subfamilies have not been thoroughly investigated, likely due to insufficient sequence information (Dittami et al., 2010). In this study as well, the diatom Lhcx subfamily and the basal streptophyte algae and core green algae Lhcsr subfamily formed a monophyletic group (Figure 1). The Lhcx/Lhcsr clade formed by these showed strong support for monophyly with the diatom Lhcq, CgLhcr9 homolog, and Lhcf subfamilies, and was in a different clade from the green algae Lhca/Lhcb subfamily, clearly indicating that it is derived from red-lineage algae. In particular, since the Lhcx/Lhcsr clade showed monophyly with the CgLhcr9 homolog and Lhcf subfamily, it was demonstrated that the Lhcx subfamily is one of the new LHC subfamily groups that arose in heterokont algae along with Lhcq, CgLhcr9 homolog, and Lhcf subfamilies. Furthermore, since Lhcsr is conserved in both core green algae and streptophyte algae, this supports the hypothesis proposed in Dittami et al. that the ancestor of extant green plants acquired the Lhcx/Lhcsr subfamily through horizontal gene transfer from a red-lineage alga, presumably a heterokont alga or haptophyte, before the last common ancestor of extant green plants. The estimated divergence time of extant green plants into Streptophyta and Chlorophyta overlaps with the emergence of heterokonts (Strassert et al., 2021), indicating that gene transfer of Lhcx/Lhcsr from heterokonts to ancestral green plants could have occurred prior to this divergence event. It is difficult to determine whether this gene transfer was merely a horizontal gene transfer or involved the acquisition of a temporary red-lineage plastid, such as a kleptoplast. Additionally, looking at Figure 1, the monophyletic clade of the green algae Lhca and Lhcb subfamilies does not form a monophyletic group with the red algae Lhcr subfamily, and between them exist cryptophyte Lhcr and diatom Lhcr. This does not negate the possibility that not only Lhcsr but also the Lhca and Lhcb subfamilies were acquired through horizontal gene transfer. This result calls for reconsideration of the formation and establishment of early chloroplasts.

Within Lhcx, Lhcx6_1, which exists at the most basal position, is present in *T. pseudonana* and *Cyclotella cryptica* but not in *C. gracilis* or *Phaeodactylum tricornutum* (Figure 1, inset). This Lhcx6_1 not only has an evolutionary distance from other typical Lhcx in the molecular phylogenetic tree but also does not show a low NPQ phenotype when deficient (Nakayasu et al., 2024). Furthermore, Lhcx6_1 directly binds to PSII and has a completely different assembly from *C. gracilis* PSII–FCPII (Feng et al., 2023; Zhao et al., 2023). Based on these observations, it is considered that Lhcx6_1 is an atypical Lhcx and does not mediate NPQ, and we argue that its classification within the Lhcx subfamily should also be reconsidered.

### Spectroscopic Characteristics of Lhcx Quenching

In this study, we successfully obtained NPQ signals at 140 ps around 687 nm by measuring time-resolved fluorescence comparing frozen whole cells immediately after dark adaptation and HL irradiation, reflecting the *in vivo* state (Figure 6, Supplemental Figure 7). By using the *lhcx1* mutant that shows a clear phenotype with almost no NPQ induction, and by optimizing the dark adaptation and HL exposure time to 30 minutes, we were able to clearly distinguish signals attributable to Lhcx-mediated NPQ.

Our results provided the first report measuring the time-resolved fluorescence of mutants with near-complete NPQ deficiency, resulting in a clear signal of NPQ. Even though several similar attempts on diatoms and *Nannochloropsis* have been made, they could not find a clear NPQ signal in this approach in the previous studies. For example, similar analyses were performed for the diatoms *C. meneghinia* and *P. tricornutum* (Miloslavina et al., 2009). A different approach was taken regarding DAS fitting compared to this study, with strong assumptions placed on the fluorescence signals of PSII and PSI, and quenching signals like those observed in this study were not obtained. More detailed observations of quenching using time-resolved fluorescence have been attempted in the diatom *C. meneghiniana*, but the ∼200 ps signal around 685 nm observed in this study was not observed; instead, a ∼600 ps quenching signal at 695 nm was observed and attributed to PSII (Chukhutsina et al., 2014). However, since they only used WT, they could not conclude the signal of ∼600 ps as NPQ depending on Lhcx. In a similar approach, also using only WT of *Nannochloropsis*, a 214 ps signal at 685 nm in *Nannochloropsis gaditana* was attributed to LHCII quenching, but it was not clear whether this was a quenching signal due to the presence of Lhcx (Chukhutsina et al., 2017). A similar approach using *lhcx* mutants has been performed in *P. tricornutum* (Taddei et al., 2018). However, the *lhcx1* knockdown mutant of *P. tricornutum* showed limited decrease in the abundance of Lhcx1 and NPQ. LL-grown *lhcx1*-knockdown mutants showed a slight decrease in NPQ, and HL-grown *lhcx1*-knockdown showed no difference in NPQ because of the expression of Lhcx3, which is an isoform of the Lhcx subfamily in *P. tricornutum*. Therefore, the observation and discussion of NPQ signal in time-resolved fluorescence was limited in contrast to this study using *lhcx1*-knockout mutants, which show minimal expression of other Lhcx isoforms (Supplemental Figure 10). Note that Buck and co-workers created *Lhcx1*-knockout mutants of *P. tricornutum*, which showed a clearer drop in NPQ in contrast to knock-down mutants; however, they did not report time-resolved fluorescence (Buck et al., 2019). Considering these things together, this study provided the first clear NPQ signal in time-resolved fluorescence analysis using almost-complete knock-out of Lhcx in Heterokonts, including diatoms.

The spectroscopic characteristics of NPQ in diatoms differ notably from those in green algae. In green alga *Chlamydomonas reinhardtii stt7-9* state-transition-less mutants, NPQ signals were observed at 150 ps in time-resolved fluorescence and ∼100 ps in transient absorption at physiological temperature, similar to our time constant, but the NPQ signal from fluorescence was detected at 682 nm and attributed to LHCII fluorescence (Tian et al., 2019; Zheng et al., 2024). This wavelength difference (682 nm vs. 687 nm) likely reflects differences in measurement temperatures and distinct pigment compositions between the two algal groups, rather than fundamental mechanistic differences.

Time-resolved fluorescence measurements in whole cells revealed longer excitation energy transfer times compared to isolated complexes. While excitation energy transfer from FCPII to the PSII core in isolated C_2_S_2_M_2_ PSII–FCPII occurs up to a time constant of 210 ps (Nagao et al., 2022), our measurements in dark-adapted whole cells showed a significantly longer time constant of 640 ps (Figure 6, Supplemental Figure 7). This threefold increase suggests that PSII–FCPII complexes *in vivo* possess larger antenna systems than the isolated C_2_S_2_M_2_ form, indicating the presence of additional FCPs connected to the excitation energy network. These peripheral antenna complexes, likely including L-FCPII dimers, extend the effective antenna size and modify energy transfer kinetics.

### The mechanism of Lhcx1-mediated NPQ

The localization of diatom Lhcx has been debated for many years. Except for Lhcx6_1 associated with PSII core (Feng et al., 2023; Zhao et al., 2023), the localization of Lhcx has been detected in free FCP or bulk FCP (Lepetit et al., 2010; Grouneva et al., 2011), except for reports that *P. tricornutum* Lhcx1/Lhcx2 bind to PSI (Grouneva et al., 2011). Additionally, in SDS-PAGE following CN-PAGE, *T. pseudonana* Lhcx1 was detected as a free monomer (Zhou et al., 2024). In the 2D-CN/SDS-PAGE of the diatom *C. gracilis* by Zhou et al. (2024), Lhcx1 was not detected. In this study, we utilized Amphipol A8-35, a surfactant that enhances the stability of membrane proteins, to enable the separation of diatom C_2_S_2_M_2_ PSII–FCPII, which had been previously challenging with CN-PAGE methods (Figure 7A). Under this condition, the dimer (L-dimer) consisting of CgLhcf12/CgLhcf2 (FCP-B/C) showed a broad band in the first dimension (Figure 7B), and a similar band pattern was observed in the immunoblot for CgLhcx1 (Figure 7C). These findings suggest that CgLhcx1 may interact with the FCP L-dimer.

Assuming the interaction between L-FCPII dimers and Lhcx1, we propose that Lhcx1-mediated NPQ operates through two possible mechanisms: either Lhcx1 directly functions as the quenching center, or quenching occurs through the L-dimer via its interaction with Lhcx1. This mechanism can be classified within the framework of Q1 and Q2 quenching, where Q1 occurs in detached antenna complexes and Q2 occurs in minor LHCs bound to the PSII core (Holzwarth et al., 2009; Miloslavina et al., 2009; Chukhutsina et al., 2014). Given that L-FCPII dimers bind loosely to S/M-FCPII, Lhcx1-mediated quenching likely represents Q1-type quenching occurring in the peripheral light-harvesting antenna, although it is unclear whether the L-FCPII dimer detaches from PSII-S/M-FCPII.

The 140 ps quenching time constant observed in HL-treated whole cells is notably faster than previously reported values for isolated FCP complexes. Isolated FCP-A tetramers and FCP-B/C dimers have been reported to show characteristic quenching with time constants of 270 ps and 200 ps, respectively (Nagao et al., 2013). This acceleration in whole cells may result from two factors: first, isolated FCP-B/C complexes lack Lhcx proteins, which appear crucial for efficient quenching; second, isolated complexes lack the native membrane environment, including the ΔpH gradient and three-dimensional protein organization necessary for optimal quenching function. These results highlight the importance of studying NPQ mechanisms in intact cellular systems rather than relying solely on isolated protein complexes.

### Compensatory mechanisms to the Lhcx1 Deficiency

The *lhcx1* strain, which lacks the most highly expressed Lhcx1 in *C. gracilis* created using genome editing, induced almost no NPQ and showed higher PSII effective quantum yield and oxygen-evolving activity compared to the WT when acclimated to HL. Since NPQ mutants in land plants show decreased PSII electron transport activity, i.e., decreased effective quantum yield, under HL acclimation (Yang et al., 2022), it is unusual that *lhcx1*, an NPQ mutant in this study, shows a higher PSII effective quantum yield. Since *lhcx1* has a smaller antenna size even when dark-adapted after HL acclimation (Figure 6, Supplemental Figure 7), the decrease in excitation energy acquisition per PSII core is thought to contribute to a higher PSII effective quantum yield. Additionally, since the PSII effective quantum yield increased even under LL conditions with the addition of CO_2_ (Supplemental Figure 9), and transcriptomic analysis revealed that *lhcx1* specifically enhances carbon fixation and central carbon metabolism pathways under both ambient air and CO_2_ conditions (Figure 8B), it is likely that the increase in carbon fixation capacity in the downstream electron transport chain also contributed to the improvement of PSII effective quantum yield. The activation of downstream electron sinks can be observed under HL conditions (Figure 2C).

HL acclimation in *lhcx1* was also suggested by the pigment analysis (Figure 3C–F). Chl content per cell decreased in *lhcx1*, and while the amount of Fx per Chl did not change, the amounts of xanthophylls Dtx and Ddx increased. The de-epoxidation state (DEPS) calculated from Dtx and Ddx also increased in *lhcx1* under HL, indicating that acclimation responses under HL are enhanced from the perspective of pigment composition as well. Generally, in WT diatoms, the amount of Dtx is very well correlated with NPQ intensity (Lavaud and Lepetit, 2013). However, since *lhcx1* strains show almost no induction of NPQ despite increased Dtx amounts, the correlation between Dtx amount and NPQ requires the presence of Lhcx. Although it is not clarified whether the increased xanthophylls in *lhcx1* are incorporated in LHC/FCP, they may function as antioxidants in thylakoid membranes and contribute to photoprotection. The above multiple photoprotection mechanisms that became apparent in the absence of Lhcx1 would contribute to the adaptation in variable marine environments.

## Materials and Methods

### Phylogenetic Analysis of LHCs Across Various Photosynthetic Eukaryotic Lineages

LHC sequences were collected from genomes and transcriptomes included in the Phycocosm and EukProt databases, in addition to our previous dataset (Kumazawa et al., 2022; Kumazawa and Ifuku, 2024), following the previous method (Kumazawa and Ifuku, 2024). The LHCs in the dataset were obtained from red algae, early-diverging streptophyte green algae (Leliaert et al., 2012): *Mesostigma viride* NIES-296 (Liang et al., 2020) and *Klebsormidium nitens* NIES-2285 (Hori et al., 2014), diatoms: *C. gracilis*, *Thalassiosira pseudonana*, *Cyclotella cryptica*, *Phaeodactylum tricornutum*, and cryptophytes: *Guillardia theta* CCMP2712 (Curtis et al., 2012) and Cryptophyceae sp. (*Baffinella* sp.) CCMP2293 (Dorrell et al., 2023). As reference sequences for Lhcsr, *Chlamydomonas reinhardtii* Lhcsr1 (UniProt: P93664) and Lhcsr3.1 (UniProt: P0DO19), which is identical to Lhcsr3.2 (UniProt: P0DO18), were added. Redundant sequences were removed from the LHC protein sequences using CD-HIT (Fu et al., 2012), and alignment was performed with mafft-einsi v7.525, followed by manual curation (Katoh and Standley, 2013). The curated dataset was realigned, and trimming was performed using ClipKit v2.3.0 in smart-gap mode to remove unaligned sites (Steenwyk et al., 2020). The phylogenetic tree was estimated using IQ-TREE v2.3.6 with the Q.pfam+I+R7 model selected by ModelFinder based on LG and Q.pfam models (Kalyaanamoorthy et al., 2017; Minh et al., 2020; Minh et al., 2021), and visualized with iTOL v6 (Letunic and Bork, 2024). Ultra-fast bootstrap approximation (UFBoot) and SH-aLRT support (%) tests were performed with 1000 replicates each (Guindon et al., 2010; Hoang et al., 2018). The aBayes support test was also conducted (Anisimova et al., 2011). The annotation of LHC subfamilies in the phylogenetic tree was based on our previous studies (Kumazawa et al., 2022; Kumazawa and Ifuku, 2024) and drawn on the tree using InkScape software.

### Cultivation

The marine centric diatom *C. gracilis* (UTEX LB 2658) was used for all analyses. Pre-cultivation was performed in IMK medium —4% (w/v) sea salt (SIGMA S9883, US) or artificial seawater made with Daigo’s Artificial Seawater SP (Nihon Pharmaceutical MS, Japan) supplemented with 1x Daigo’s IMK medium (Nihon Pharmaceutical MS, Japan) and 0.2 mmol L^−1^ Na_2_SiO₃— at 20°C under fluorescent lights at approximately 30 µmol photons m^−2^ s^−1^ with continuous shaking at 100 rpm. Seven–eight-day or three-day cultivation was performed in IMK medium at 25°C under warm white LEDs at 30 µmol photons m^−2^ s^−1^ (LL) or 300 µmol photons m^−2^ s^−1^ (HL), with continuous aeration of air or 3% (v/v) CO_2_.

### Preparation of Thylakoid Membranes from *C. gracilis* and Spinach

Diatom WT cells cultured under HL at 25°C, 300 µmol photons m^−2^ s^−1^ for three days were centrifuged at 1,200×*g*, 20°C for 5 minutes to collect the cells, then resuspended in artificial seawater, washed, and centrifuged again. After removing the supernatant, the pellet was resuspended in 1 mL of artificial seawater, transferred to a 1.5 mL microcentrifuge tube, and centrifuged at 21,500×*g*, 4°C for 5 minutes to obtain a pellet. The pellet was resuspended in 1 mL of cell disruption buffer (20 mM Tricine-KOH (pH 7.6), 0.4 mol L^−1^ sorbitol, 10 (w/v) % PEG-6000, 10 mM EDTA), mixed with 0.5 g of 0.1 mm diameter glass beads, and processed with a bead beater for 10 seconds. The sample was then well-chilled on ice, and the disruption process was repeated a total of three times. The supernatant was centrifuged at 21,500×*g*, 4°C for 5 minutes to obtain a pellet. The pellet was resuspended in 1 mL of washing buffer (20 mmol L^−1^ Tricine-KOH (pH 7.6), 0.4 mol L^−1^ Sorbitol, 5 mmol L^−1^ MgCl_2_, 2.5 mmol L^−1^ EDTA) and centrifuged at 50×*g*, 4°C for 5 minutes. The portion containing thylakoid membranes was carefully collected, avoiding the upper and lower contaminants, and centrifuged at 21,500×*g*, 4°C for 5 minutes to obtain a thylakoid membrane pellet. The pellet was resuspended in solubilization buffer (50 m mol L^−1^ imidazole-HCl (pH 7.0 at 4℃), 20 (v/v) % glycerol), and a portion was taken to measure the concentrations of Chl *a* and *c* (Jeffrey and Humphrey, 1975). The thylakoid membrane of spinach (*Spinacia oleracea*) was isolated from market spinach based on previously reported methods (Yamamoto et al., 2011) using the modified buffer (50 mmol L^−1^ tricine-NaOH (pH 7.8 at room temperature), 0.4 L^−1^ sucrose, 10 mmol L^−1^ NaCl, 5 mmol L^−1^ MgCl_2_). The thylakoid membranes were adjusted to 1 μg μL^−1^ of Chl *a*+*c* with solubilization buffer (50 mM imidazole-HCl (pH 7.0 at 4℃), 20 (v/v) % glycerol).

### 2D-CN/SDS-PAGE

The 4–13% gradient gel used for CN-PAGE was prepared according to existing methods (Järvi et al., 2011; Kameo et al., 2021), and the entire CN-PAGE procedure followed the modified CN-PAGE method using Amphipol A8-35 (Kameo et al., 2021). However, 75 mmol L^−1^ imidazole-HCl (pH 7.0 at 4°C) was used in the 3x gel buffer, and 40 (w/v) % acrylamide/bis solution (37.5:1) (019-25655, Fujifilm Wako, Japan) was used for acrylamide. The anode buffer contained 25 mmol L^−1^ imidazole-HCl (pH 7.0 at 4°C), and the cathode buffer contained 50 mmol L^−1^ Tricine, 7.5 mmol L^−1^ imidazole. The Chl concentrations in resuspended thylakoids were measured (Jeffrey and Humphrey, 1975), and thylakoid membranes were adjusted to 1 μg μL^−1^ of Chl *a*+*c* and Chl *a*+*b* with solubilization buffer (50 mmol L^−1^ imidazole-HCl (pH 7.0 at 4℃), 20 (v/v) % glycerol) and resuspended. An equal volume of 2 (w/v) % dodecyl-α-D-maltoside (α-DM, Anatrace, OH, USA) was added for solubilization, placed on ice for 2 minutes, then centrifuged at 21,500×*g* for 2 minutes to remove insoluble residues. The supernatant was mixed with half the volume of 2 (w/v) % amphipol A8-35 (Anatrace, OH, USA), and 5 μg of Chl in 15 μL was applied to each well. Electrophoresis was performed using MyPowerII 300 power supply (AE-8135, ATTO, Japan) at 40 V, 1 mA for 660 minutes at 4°C with two mini gels simultaneously. After electrophoresis, the gels were scanned with the scanner ES2200 (EPSON, Japan). The CN-PAGE gels were immersed in 50 (v/v) % glycerol on ice, cut into gel strips, and stored at − 80°C in 50 (v/v) % glycerol until use.

Gel strips were solubilized by immersing them in solubilization buffer (1% SDS and 50 mM dithiothreitol (DTT) for 30 minutes at room temperature, and then applied to gels with a stacking gel (pH 6.8) and separation gel (14% acrylamide (37.5:1), 6 mol L^−1^ Urea, pH 8.6). The Laemmli system was used for electrophoresis. When the dye had electrophoresed to the bottom of the marker well at 250 V, 100 mA, the process was temporarily stopped, and Precision Plus Protein Dual Color Standards (#1610374, BioRad, CA, USA) were applied. Electrophoresis was then continued at 250 V for 90 minutes. After electrophoresis, gels not used for immunoblotting were fluorescently stained with Oriole fluorescent gel stain (#1610496, BioRad, CA, USA) for 100 minutes, gently rinsed with distilled water, and then fluorescence was detected using the ChemiDoc Touch Imaging System (Bio-Rad).

### Construction of hCas9 Vector

The hCas9 sequence was amplified from the hCas9 vector (Addgene, plasmid #41815) using the InF-hCas9-fw1 primer (TTTATAAAGCGGATCATGGACAAGAAGTACTCCATTGG) and InF-hCas9-rv1 primer (AACAGCTTGCATGCCTCACACCTTCCTCTTCTTCTTG). The PCR product was cloned into the linearized pCgNRp vector digested with BamHI and PstI using In-Fusion reaction according to the manufacturer’s protocol. The obtained clone vector was named pCgNRphCas9.

### Construction of pCgNRpble-Lhcx1-RGR Vector

A 20-mer genome editing target sequence containing the PAM (protospacer adjacent motif) sequence (NGG) for gRNA in the CRISPR/Cas9 system, GTTGCTATGCTTGCCGTTAT**TGG** (bold indicates the PAM sequence), was selected. A system that expresses complete gRNA in cells via a mRNA-type Pol II promoter by connecting self-processing ribozymes (Gao and Zhao, 2014; Zhang et al., 2017) was used. This method was adapted to the genome editing system of *C. gracilis*. The genome editing vector containing gRNA was constructed using three primers: Lhcx1-RGR-F1; GACGAAACGAGTAAGCTCGTC**GTTGCTATGCTTGCCGTTAT**GTTTTAGAGCTAGAAATAGCAAG (bold indicates the 20-mer sequence for gRNA target recognition excluding the PAM sequence), Lhcx1-RGR-F2; GTCA<u>GGATCC</u>**AGCAAC**CTGATGAGTCCGTGAGGACGAAACGAGTAAGCTCGTC (BamHI restriction enzyme site is underlined, and bold indicates the complementary sequence of the first six bases of the 20-mer for gRNA target recognition including the PAM sequence), and PstI-RGR-Rv; ATAT<u>CTGCAG</u>GTCCCATTCGCCATGCCGAAGC (PstI restriction enzyme site is underlined). The PstI-RGR-Rv primer is a universal primer in this protocol. The first fragment (199 bp) was amplified by PCR from pRS316-RGR-GFP (Addgene, plasmid #51056) using the Lhcx1-RGR-F1 and PstI-RGR-Rv primers. The second fragment (231 bp) was amplified by PCR from the first purified PCR product using the Lhcx1-RGR-F2 and PstI-RGR-Rv primers. The purified second product was digested with BamHI (NEB) and PstI (NEB), and ligated with the pCgNRpble vector linearized with BamHI and PstI. The final product was named pCgNRp^ble^-Lhcx1-RGR.

### Transformation

The electroporation transformation procedure using the Super electroporator NEPA21 Type-II (NEPAGENE, Japan) followed previous reports (Miyahara et al., 2013; Ifuku et al., 2015; Ifuku and Yan, 2020). The initial transformation of the *hCas9* gene into *C. gracilis* was performed using the pCgNRphCas9 vector linearized with DraIII and SapI. Colonies were selected on IMK plates containing the antibiotic Nourseothricin at 400 µg mL^−1^ (clonNAT; Hans Knoell Institute, Jena, Germany). The introduced *hCas9* gene was confirmed by PCR using the first set: InF-hCas9-fw1 primer and hCas9-2088-Rv primer: CCTTAAAGGTGAGAGAGTCATCATGGATC, and another set: hCas9-1965n-Fw primer: ATGAAACAGCTCAAGAGGCGCCGA and InF-hCas9-rv1 primer. The hCas9 strain (hC9 strain) used for further genome editing was selected from eight transformants screened by PCR and immunoblotting using anti-Cas9 antibody (described later).

The linearized pCgNRpble-Lhcx1-RGR was introduced into the hC9 strain by electroporation. This was essentially the same as the transformation of hCas9. Before introducing the RGR vector, the vector was linearized with HindIII (NEB) and concentrated to 850–1000 ng µL^−1^ by ethanol precipitation. The amount of linearized vector was 5 µg per electroporation. Colonies were selected on IMK plates containing the antibiotic Zeocin at 100 µg mL^−1^ (InvivoGen #ant-an-1, US CA). Mutations were confirmed by Sanger sequencing of the products amplified by PCR using Lhcx1_seq_FW primer: AGAGACTGTAAGGATTCGGA and Lhcx1_seq_RV primer: GAAGGAAGATCCCTCTACCT.

### Immunoblotting

Diatom cells were collected and solubilized in sample buffer containing 62.5 mmol L^−1^ Tris–HCl (pH 6.8), 2.5% (w/v) SDS, 10% (w/v) glycerol, 2.5% (v/v) 2-mercaptoethanol, and a trace amount of bromophenol blue (BPB). Solubilization was performed by incubating at 98°C for 6 minutes, followed by 30 minutes at room temperature, and then centrifugation. The protein concentration was adjusted to 30 μg/20 μL. Thirty micrograms of protein was applied to each lane of an 8% acrylamide separation gel with a 3% acrylamide stacking gel and separated by SDS-PAGE. Proteins were transferred to an Immobilon-P PVDF membrane (Millipore, MA, USA) and incubated with anti-Cas9 polyclonal antibody (632607, Takara, Japan) at a 1:2000 dilution.

For the detection of Lhcx1, SDS-PAGE gels were prepared with a 15% acrylamide separation gel and a 4.5% acrylamide stacking gel. Two μg of protein was applied to each lane and separated by SDS-PAGE. The anti-Lhcx1 peptide antibody was produced against the synthetic peptide “KEEYAPGDLRFDPFGLMP”. After electrophoresis, the gel was desalted and equilibrated in Electron transfer buffer (25 mmol L^−1^ Tris, 192 mmol L^−1^ glycine, 20 (v/v) % methanol, 0.05 (w/v) % SDS). Subsequently, proteins were transferred from the gel to an Immobilon-P PVDF membrane (Millipore, Billerica, MA, USA) and incubated with anti-Lhcx1 antibody at a 1:5000 dilution. Immunoreactive proteins were detected using the Amersham ECL Prime Western Blotting Detection Reagent (Cytiva RPN2236) with the ChemiDoc Touch Imaging System (Bio-Rad).

### PAM Fluorescence Measurement

Chl fluorescence by pulse-amplitude modulation was detected using an actinic light of 450 or 455 nm (blue) at 300 µmol photons m^−2^ s^−1^ with the NPQ2 protocol (200 seconds of light period and 390 seconds of dark recovery, with the first saturating pulse light in the light period at 10 seconds after the start of illumination) of AquaPen AP-100 or AP-110 (PSI, Czech Republic) using a bandpass filter (667 to 750 nm). The measuring pulse was set to 10%, and the super pulse (saturating pulse light) was set to 30% (equivalent to 900 µmol photons m^−2^ s^−1^). Cells used in the NPQ2 protocol were dark-adapted for more than 10 minutes before measurement.

### Oxygen-Evolving Activity Measurements

The diatom cells grown under HL conditions (200 μmol photons m^−2^ s^−1^ with rotational shaking at ∼22°C) were adjusted to a concentration of 5 μg Chl *a* mL^−1^ (Jeffrey and Humphrey, 1975), following the methods described in Nakayasu et al. (Nakayasu et al., 2024). Using a Clark-type oxygen electrode (Hansatech, UK) at 25°C, oxygen evolution was measured in the medium supplemented with 10 mM NaHCO_3_. Cells were illuminated with white light at 300 μmol photons m^−2^ s^−1^, and the cell suspension in the reaction chamber was continuously stirred with a magnetic stirrer during measurements.

### HPLC Pigment Analysis

Cells cultured for 7–8 days under LL or HL conditions were collected by centrifugation at 1,500 x *g* for 5 minutes, and the supernatant was removed. The pellet was immediately frozen in liquid nitrogen and stored at −80 °C until extraction. The frozen pellet was extracted using 100% acetone (Wako, Japan) with sonication for 6 minutes in an ice-water bath. Subsequently, the sample was centrifuged at 15,000 rpm for 5 minutes at 4 °C. The resulting supernatant was filtered through a 0.45 nm hydrophilic PTFE filter (SLPT0445NL, Hawach Scientific, IL, USA) and subjected to HPLC. The HPLC method is based on Nagao, Ueno, et al. (2020) with modifications derived from Manuel Zapata et al. (2000). The 0.25 mol L^−1^ pyridine contained in solution A was adjusted to pH 5.0 using acetic acid. A Shimadzu HPLC system equipped with LC-20AD pumps and an SPD-M20A detector was used for pigment composition analysis. The system was equipped with an Inertsil C8 reverse-phase column (5020-01228, GL Sciences, Japan). The sample injection volume was 20 μL for each analysis. Standard pigments of Chl *a*, fucoxanthin, diadinoxanthin, and diatoxanthin were purchased from DHI (Denmark), analyzed similarly, and used to create calibration curves.

### 77K Steady-State Fluorescence Spectra

Cells for measurement were cultured under HL at 300 μmol photons m^−2^ s^−1^ at 25°C for 3 days. Samples were diluted with IMK medium and adjusted to 10 μg Chl *a*+*c*/150 μL. Chl concentration was measured after extraction with 90% acetone (Jeffrey and Humphrey, 1975). The diluted samples were treated in the dark or under HL at 300 μmol photons m^−2^ s^−1^ for 30 minutes and immediately frozen in liquid nitrogen. Low-temperature steady-state fluorescence emission spectra of whole cells were measured at 77K with excitation at 459 nm using a JASCO FP-8500 equipped with a PMU-130 liquid nitrogen cooling unit. The spectra were integrated from 5 consecutive measurements with a 2.5 nm sampling pitch. The spectra were normalized at the PSII CP47 peak (693–695 nm) (Andrizhiyevskaya et al., 2005; Nagao et al., 2010).

### Time-Resolved Fluorescence and Fluorescence Decay-Associated Spectra

Samples were prepared for time-resolved fluorescence (TRF) spectral measurements using the same method as for steady-state measurements. TRF spectra were measured using a time-correlated single-photon counting system in the range of 670 nm to 750 nm with wavelength intervals of 1 nm and time intervals of 18.3 ps (Hamada et al., 2017). A picosecond pulsed diode laser (PiL047X; Advanced Laser Diode Systems) operating at 459 nm with a repetition frequency of 3 MHz was used as the excitation light source to excite Chl *c* and fucoxanthin, which are the main light-harvesting pigments in diatoms. Detailed conditions for TRF measurements were described in Nagao et al. (2020). Fluorescence decay-associated spectrum (spectra) (FDAS) was constructed by global analysis (Ye et al., 2025). The analysis of fluorescence decay *I*_F_(λ, t) at wavelength λ with the instrument response function *I*_IRF_(t) followed the function:

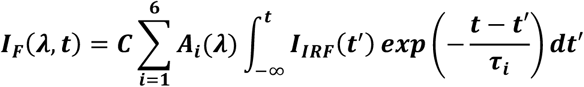

The time constants were set to τ_1–6_ = 40 ps, 140 ps, 640 ps, 1.6 ns, 3.8 ns, and 26 ns based on the analysis results of WT cells cultured under white LED light. *A*_i_(λ) represents each FDAS. The rise and decay components of fluorescence are indicated by negative and positive peaks, respectively. Positive and negative amplitudes in the same FDAS reflect energy transfer from pigments with positive amplitude to pigments with negative amplitude (Yokono et al., 2008; Akimoto et al., 2012). Positive amplitude without accompanying negative amplitude signifies energy quenching. For Figure 6 and Supplemental Figure 7, FDAS of each sample was normalized as,

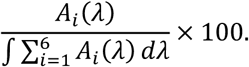

For easy viewing, a magnification factor 100 was used.

### Measurement of PSII Antenna Size

Chl fluorescence induction curves of wild-type (WT) and three *lhcx1* mutants (*lhcx1-7*, *lhcx1-15*, *lhcx1-234*) cultured under HL at 300 μmol photons m^−2^ s^−1^ at 25°C for 3 days were measured using a Fluorometer FL 3500 (PSI, Czech Republic) in the presence of 40 μM DCMU after 10 minutes of dark adaptation (Nedbal et al., 1999). The cultures were grown in two independent batches. In the measurement, Flash intensity was set to 50%, and Flash duration was set to 100 μs. The relative antenna size was determined by measuring the initial slope of fluorescence rise during the first 1 μs relative to the maximum fluorescence intensity. Four replicate measurements were performed for each strain.

For data analysis, R statistical software (R version 4.4.0 (2024-04-24)) was used, utilizing the tidyverse, readr, lme4, emmeans, and ggplot2 packages. To account for batch-specific variations, antenna size values were normalized by the WT mean value within each batch. A linear mixed model (LMM) was applied to the normalized data. Estimated marginal means were calculated, and multiple comparisons were performed using the Tukey method. The threshold for statistical significance was set at adjusted *p* < 0.05.

### RNA sequencing

Cells were harvested during the exponential growth phase after three days of treatment exposure to LL/HL and ambient/3%-CO_2_-supplemented air aeration, frozen with liquid nitrogen and stored at − 80°C until use. The frozen cells were crushed by bead-beating with 3-mm diameter zirconia beads. Total RNA was extracted using the RNeasy Plant Mini Kit (Qiagen) following the Robertson laboratory’s protocol (https://wordpress.clarku.edu/debrobertson/laboratory-protocols/rna-extraction-from-diatoms-using-the-rneasy-kit/). RNA-seq libraries were constructed using the NEBNext Ultra II RNA Library Prep Kit for Illumina (Cat No. 7770) according to the manufacturer’s instructions. Paired-end sequencing (2 × 150 bp) was performed on an Illumina NovaSeq 6000 platform. Adapter sequences (Read 1: AGATCGGAAGAGCACACGTCTGAACTCCAGTCA, Read 2: AGATCGGAAGAGCGTCGTGTAGGGAAAGAGTGT) were trimmed from raw sequencing reads. Prior to mapping, the gene structure annotation provided in ChaetoBase v1.1 (predicted_all.gff3) was modified using the following procedures: (1) replacement of geneID attributes with Parent attributes for transcripts lacking gene correspondence, (2) correction of transcripts incorrectly defined as genes, and (3) addition of missing gene definitions using AGAT v1.0.0.

Clean reads were aligned to the *C. gracilis* reference genome (ChaetoBase v1.1) using STAR v2.7.11b with the following parameters: --alignIntronMax 10000. Gene expression levels were quantified using RSEM v1.3.3 with --strandedness reverse to account for Illumina stranded RNA-seq protocol, and expected count matrices were generated for downstream analysis.

### Differential Gene Expression Analysis

Statistical analysis of gene expression was performed using the edgeR package (v4.6.2) in R (v4.5.0). Genes with low expression levels were filtered out by retaining only those with CPM (counts per million) values exceeding the 20th percentile in at least one condition. A DGEList object was created, and normalization factors were calculated using the TMM (trimmed mean of M-values) method. Dispersion parameters were estimated using the estimateDisp function, and generalized linear models (GLM) were fitted using glmQLFit.

### Statistical Contrasts and Hypothesis Testing

To address the research objectives, the following statistical contrasts were performed:

1) Primary Analysis: HL Response Comparison

WT HL response: Air.HL.WT vs Air.LL.WT

lhcx1 HL response: Air.HL.lhcx1 vs Air.LL.lhcx1

Enhanced HL response in lhcx1: (Air.HL.lhcx1 - Air.LL.lhcx1) - (Air.HL.WT - Air.LL.WT)

2) Secondary Analysis: CO_2_ Effect Validation

WT HL response under CO_2_: CO_2_.HL.WT vs CO_2_.LL.WT

lhcx1 HL response under CO_2_: CO_2_.HL.lhcx1 vs CO_2_.LL.lhcx1

Enhanced HL response in *lhcx1* under CO2: (CO2.HL.lhcx1 - CO_2_.LL.lhcx1) - (CO_2_.HL.WT - CO_2_.LL.WT)

Differential expression was determined using quasi-likelihood F-tests implemented in glmQLFTest. P-values were adjusted for multiple testing using the Benjamini-Hochberg method, with an FDR threshold of 0.05.

### Functional Enrichment Analysis

KEGG pathway enrichment analysis was conducted using the enrichKEGG function from the clusterProfiler package (v4.16.0). KEGG Orthology (KO) annotations for *C. gracilis* genes were obtained from BLASTP searches against the KEGG database using BLASTKOALA. Only pathways with q-value < 0.05 were considered significantly enriched.

## Acknowledgment

This work was supported in part by JSPS KAKENHI grant nos. JP22KJ2017 (M.K.), JP24H02086 (S.A.), JP23KJ1361 (Ko.I.), JP23H02347 (Ke.I. and S.A.), JP24H02081 (Ke.I.), and a grant from the Institute for Fermentation, Osaka, Japan (Ke.I.). We would like to thank Ms. Dongyi Yan for her help in the establishment of knock-out mutants.

## Data Availability

RNA-seq data in this study were deposited with link to BioProject accession number PRJDB12222 in the NCBI BioProject database, and the list of the data in Supplemental Data 1.

## Author Contributions

M.K. and Ke.I. conceived the project; M.K. performed phylogenetic analysis, 2D-CN/SDS-PAGE, growth experiments, PAM analyses, absorbance spectrum analyses, 77K emission fluorescence analysis, antenna size measurements, and RNA-seq *in silico* analysis, and prepared samples for ime-resolved fluorescence analyses; S.A. performed time-resolved fluorescence analyses; A.T. conceived the methodology of 2D-CN/SDS-PAGE; Ko.I. measured and analyzed oxygen-evolving activity; S.T. measured pigments; H.H., A.S., and S.I. performed RNA-seq and provided funding; N.I. conceived methodology of SDS-PAGE and immunoblotting; N.I.-K. and Y.K. provided anti-Lhcx1 antibody. M.K. drafted the original manuscript, and Ke.I. revised and edited the final manuscript. All authors contributed to the discussion of the results and improvement of the manuscript.

## Supplemental Figures and Data

**Supplemental Data 1.** Comprehensive RNA-Seq gene expression data including normalized expression values (TPM, FPKM), differential expression statistics (Log2FC, p-value, FDR), and gene annotations for all detected genes across four experimental conditions.

