## Supplemental Figure for "Evolutionary origin and functional mechanism of Lhcx in the diatom photoprotection"

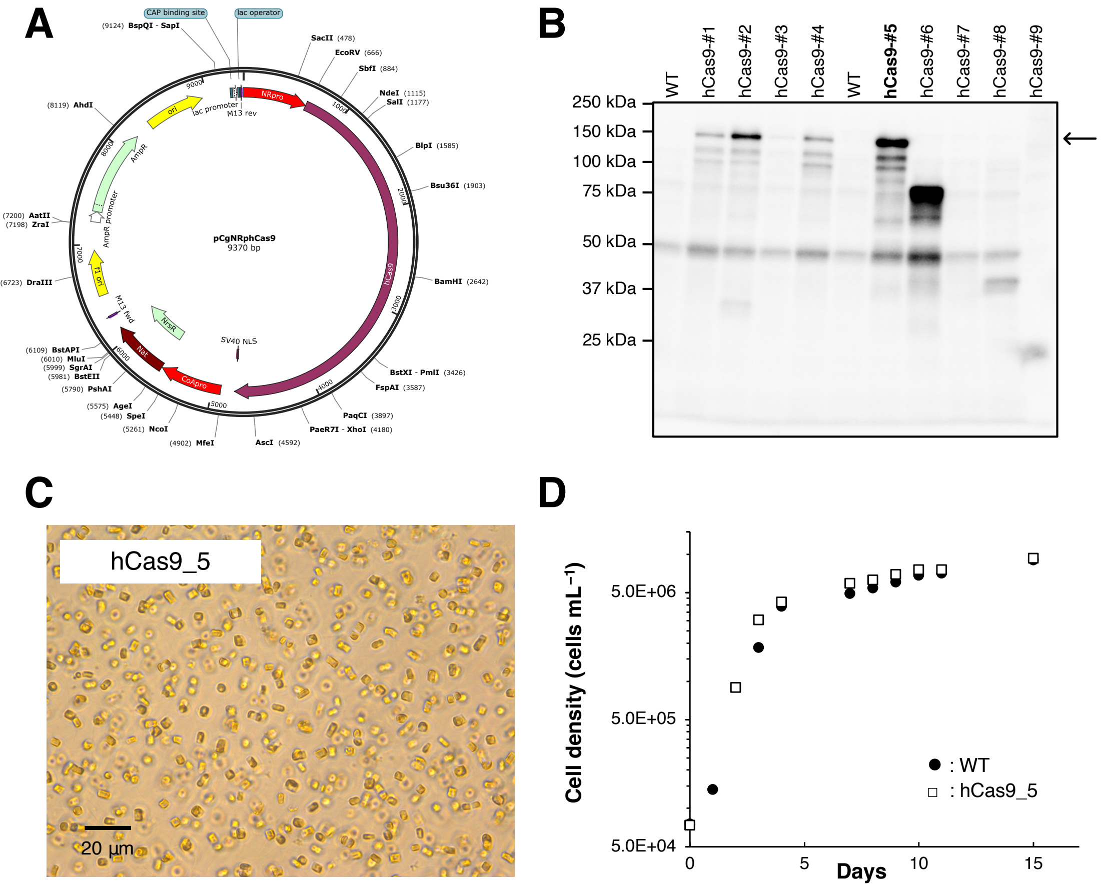


**Supplemental Figure 1. Generation of hCas9-Expressing Host Strain**

A) Plasmid map of pCgNRphCas9 for the transformation of the *hCas9* gene into *C. gracilis*. B) Immunoblot of hCas9. An arrow indicates the full-length hCas9. C) Microscopic image of the hCas9-#5 strain. D) Growth curves of wild-type (WT) and hCas9-#5 strain.

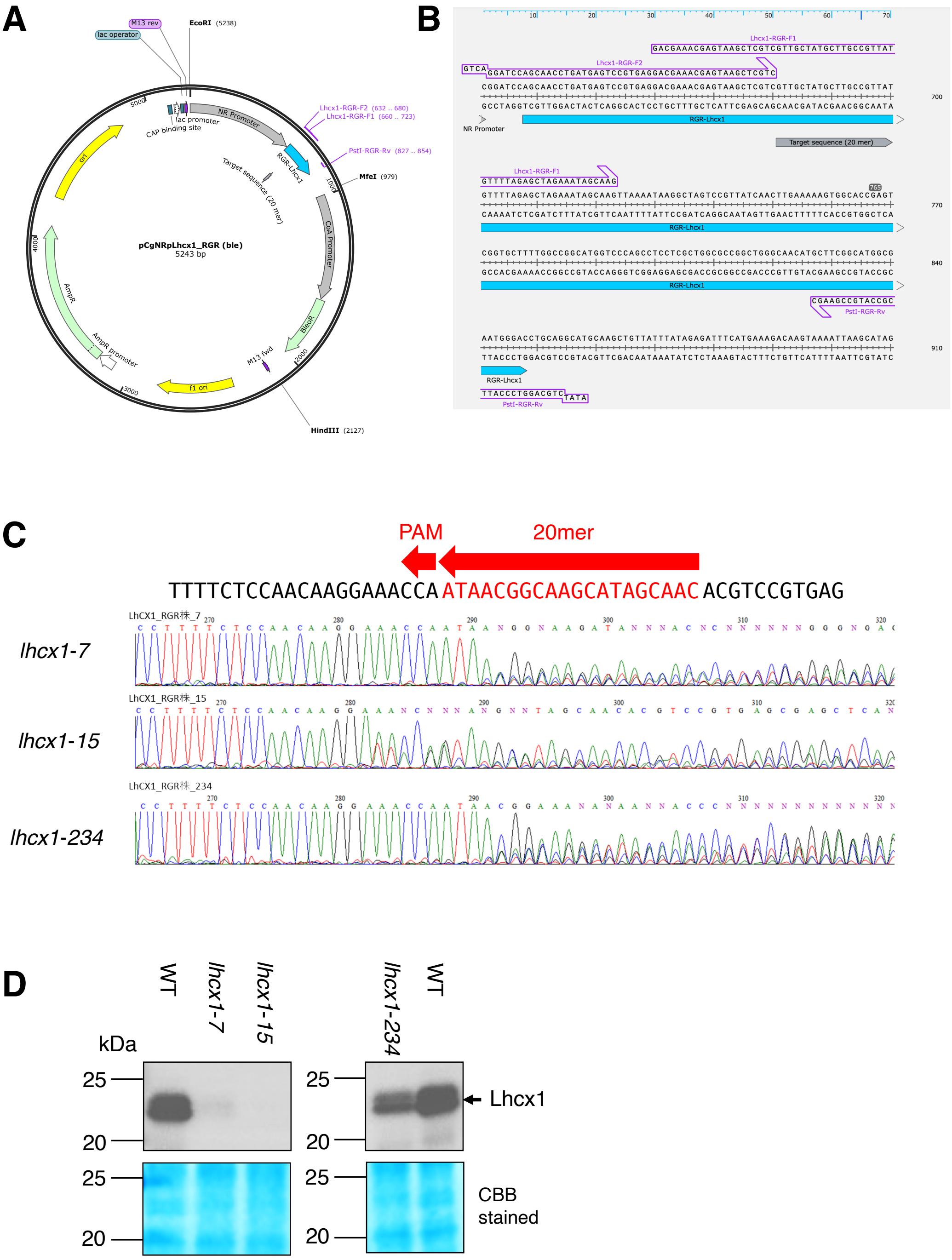


**Supplemental Figure 2. Confirmation of Lhcx1 Deficiency in *lhcx1* Genome-Edited Strains**

A) Plasmid map of the pCgNRpLhcx1-RGR for genome editing. B) The Lhcx1-RGR region of pCgNRpLhcx1-RGR. C) Sanger sequencing of the *Lhcx1* region. D) Immunoblot using anti-Lhcx1 antibody. It should be noted that there is a shift in molecular weight markers compared to Figure 7B, C, particularly in the lower molecular weight range, likely due to the composition of the SDS-PAGE gel.

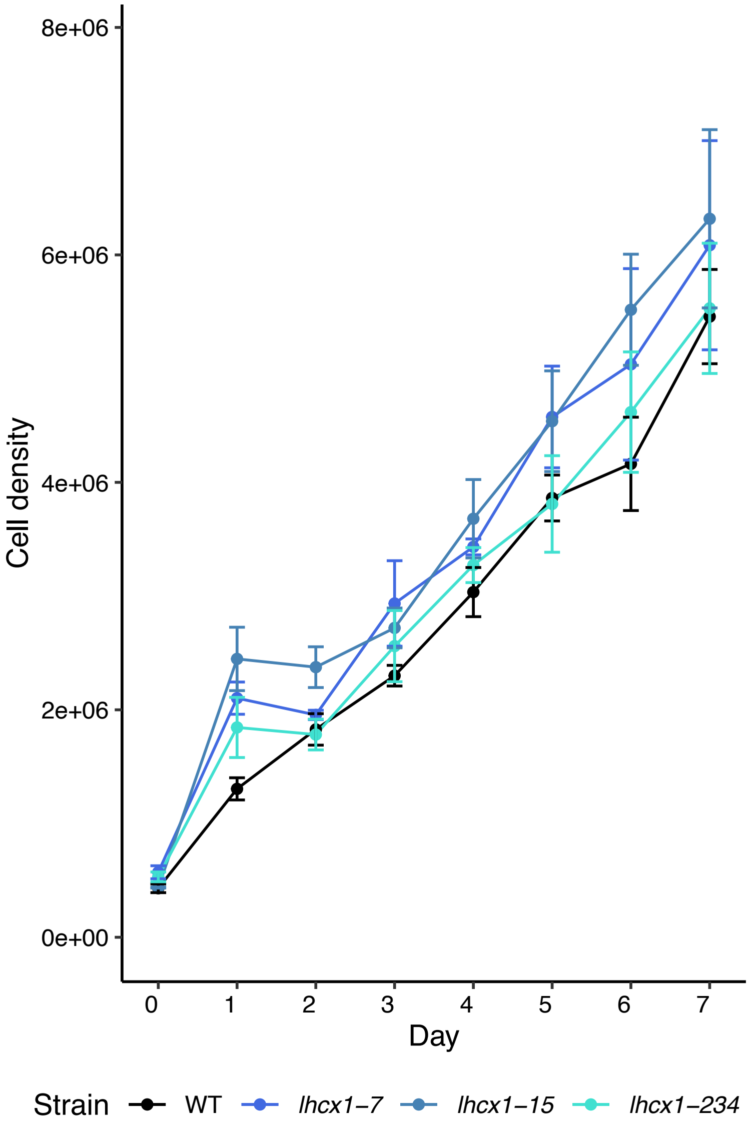


**Supplemental Figure 3. Growth Curves of *C. gracilis* Wild-Type and *lhcx1* Strains Cultured with Aeration for 7 Days Under High Light**

The points in each figure represent mean ± standard deviation with biological replicates of $n = 3$.

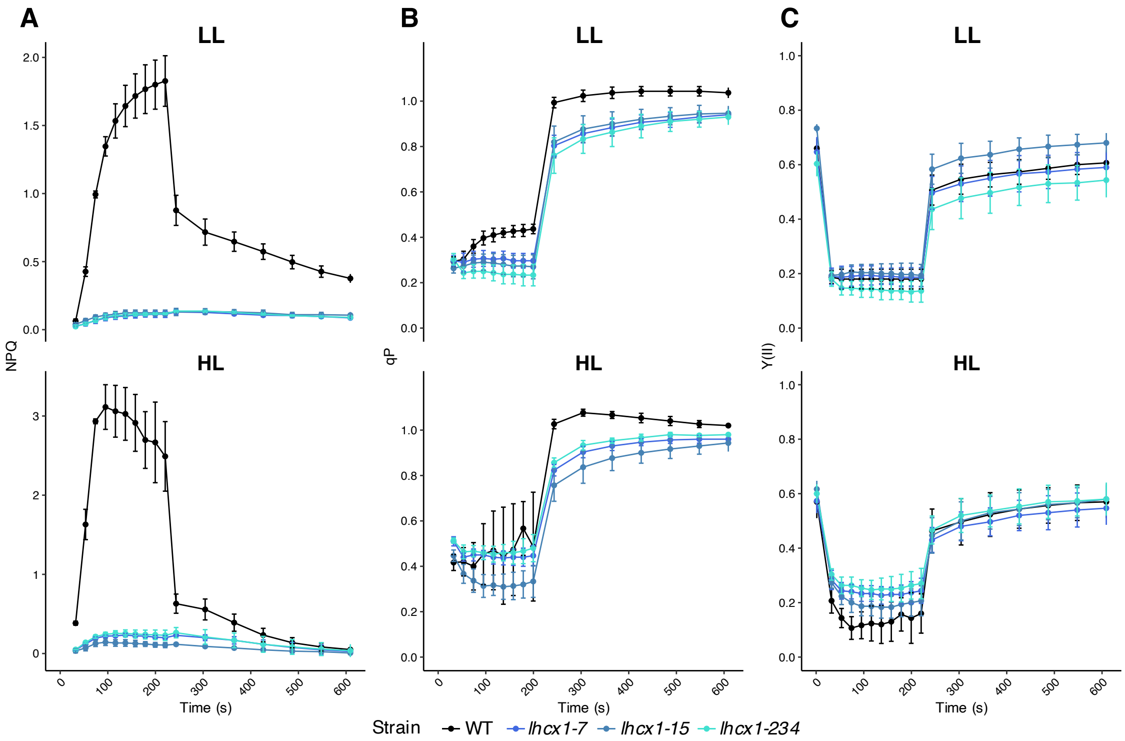


30 **Supplemental Figure 4. PAM Fluorescence Measurements of *C. gracilis* Wild-Type and *lhcx1***

31 **Strains Cultured with Aeration for 3 Days Under Low Light (LL) and High Light (HL)**

32 A) NPQ of LL and HL cultures. B) Photochemical quenching qP of LL and HL cultures. C) PSII

33 effective quantum yield Y(II) of LL and HL cultures. The points in each figure represent mean ±

34 standard deviation. with biological replicates of $n = 3$.

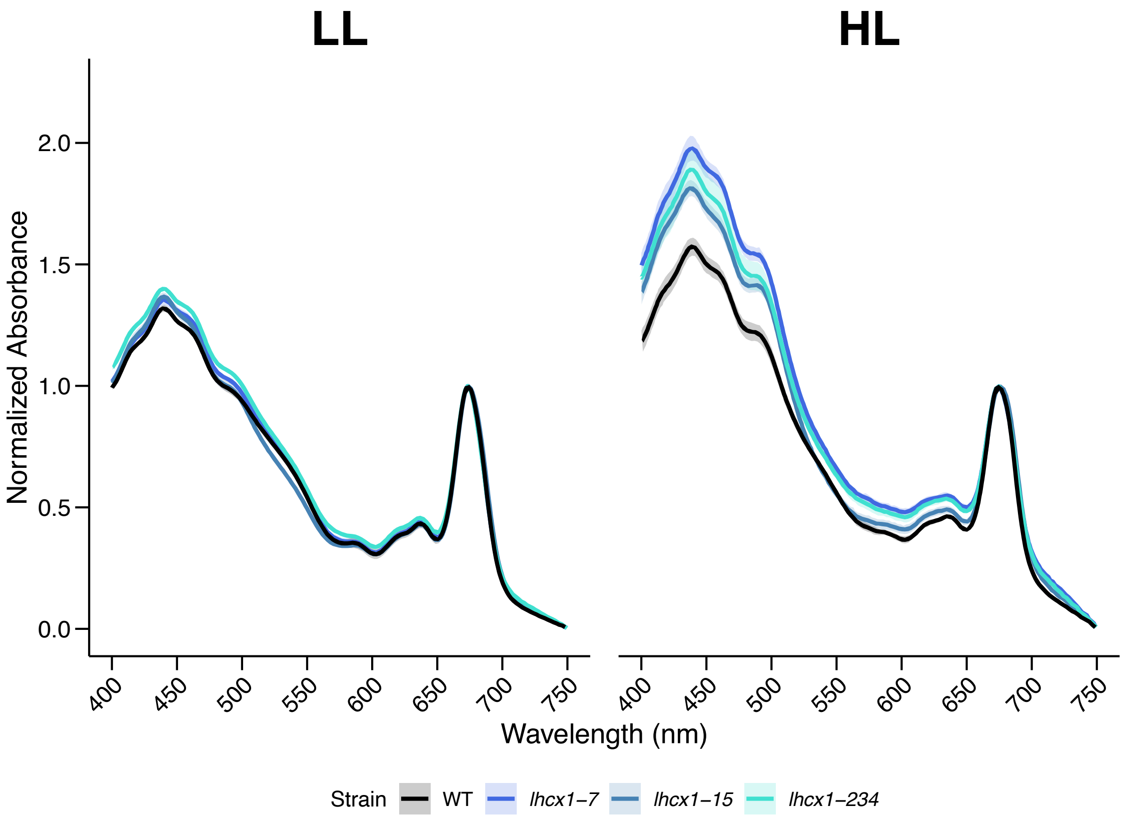


**Supplemental Figure 5. Total Absorption Spectra of Low Light (LL, left) and High Light (HL, right) Cultures**

The value at 750 nm is set to 0, and the spectra are normalized to 1 at the Qy peak of Chl *a*. The solid lines represent the mean values, and the semi-transparent areas represent the mean ± standard deviation.

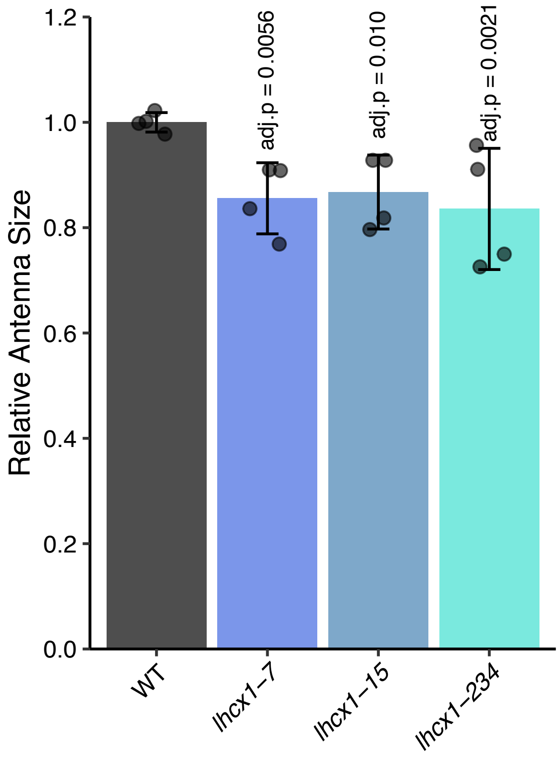


**Supplemental Figure 6. Relative PSII Antenna Size of Wild-Type (WT) and *lhcx1* Mutants**

Wild-type (WT) and three *lhcx1* mutants (*lhcx1-7*, *lhcx1-15*, *lhcx1-234*) were cultured in two independent batches. Antenna size was determined by measuring the initial slope of fluorescence rise during the first 1 μs relative to the maximum fluorescence intensity. Four replicate measurements were performed for each strain.

For data analysis, R statistical software (R version 4.4.0 (2024-04-24)) was used, utilizing the tidyverse, readr, lme4, emmeans, and ggplot2 packages. To account for batch-specific variations, antenna size values were normalized by the WT mean value within each batch. A linear mixed model (LMM) was applied to the normalized data. Estimated marginal means were calculated, and multiple comparisons were performed using the Tukey method. The threshold for statistical significance was set at adjusted $p < 0.05$.

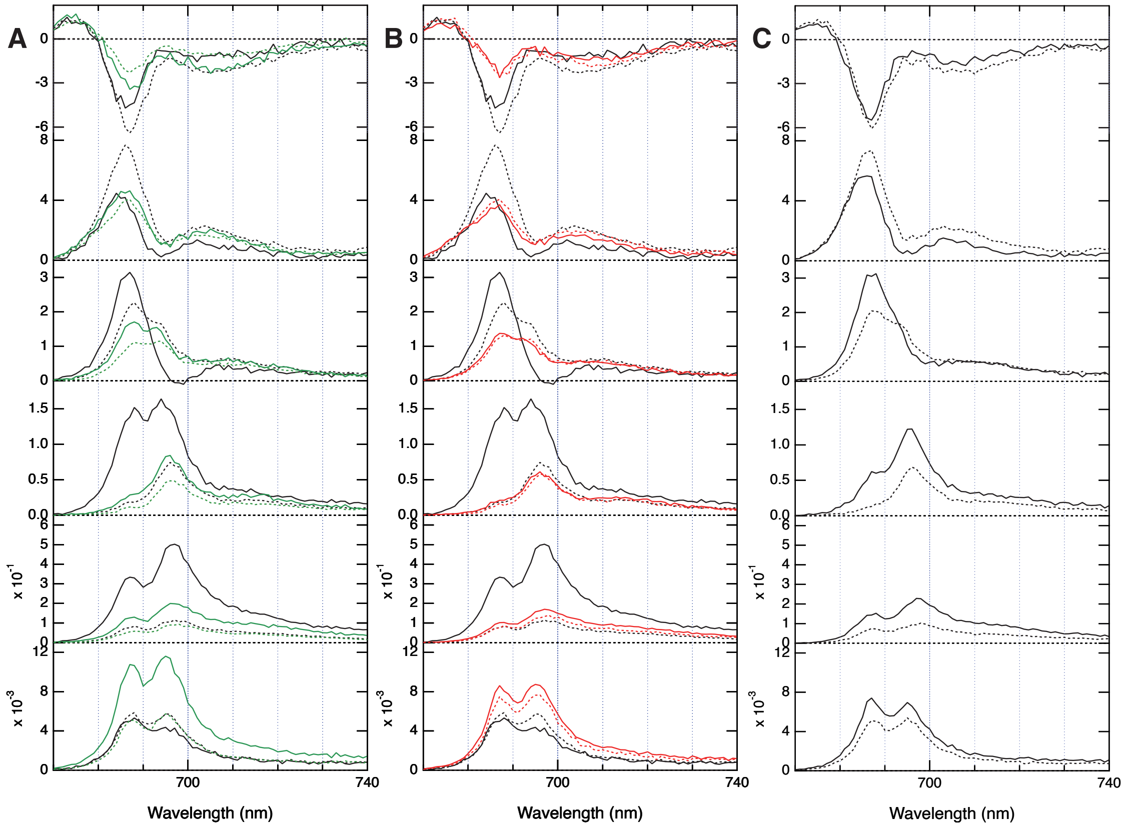


**Supplemental Figure 7. Fluorescence Decay-Associated Spectra of Time-Resolved Fluorescence of *lhcx1-15*, *lhcx1-234*, and the second replicate of WT**

Black lines indicate WT (same data as Figure 5), and green and red lines indicate *lhcx1-15* (A) and *lhcx1-234* (B). (C) The second replicate of WT. Solid lines represent measurements after 30 minutes of dark treatment, and dashed lines represent measurements after 30 minutes of treatment at 300 μmol photons $m^{-2}$ $s^{-1}$. Vertical axes are normalized fluorescence amplitude, and horizontal axes are wavelength. First to sixth boxes indicate time components of 40 ps, 140 ps, 640 ps, 1.6 ns, 3.8 ns, and 26 ns, respectively, as well as Figure 5.

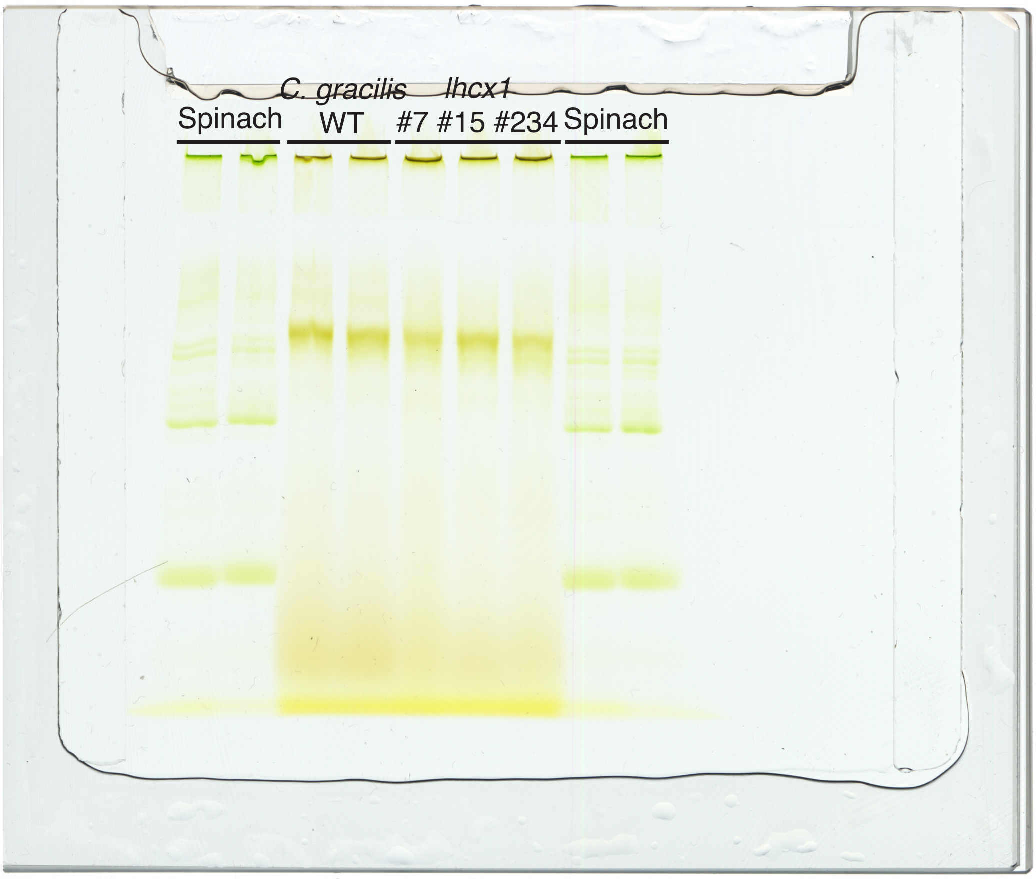


**Supplemental Figure 8. Amphipol-based-CN-PAGE of Thylakoid Membranes of *C. gracilis* WT and *lhcx1* Mutants**

Amphipol-based CN-PAGE of *C. gracilis* and spinach thylakoid membranes. Five µg Chl *a*+*c* or *a*+*b* of thylakoid membrane was solubilized with α-DDM, stabilized with amphipol, and applied to each well.

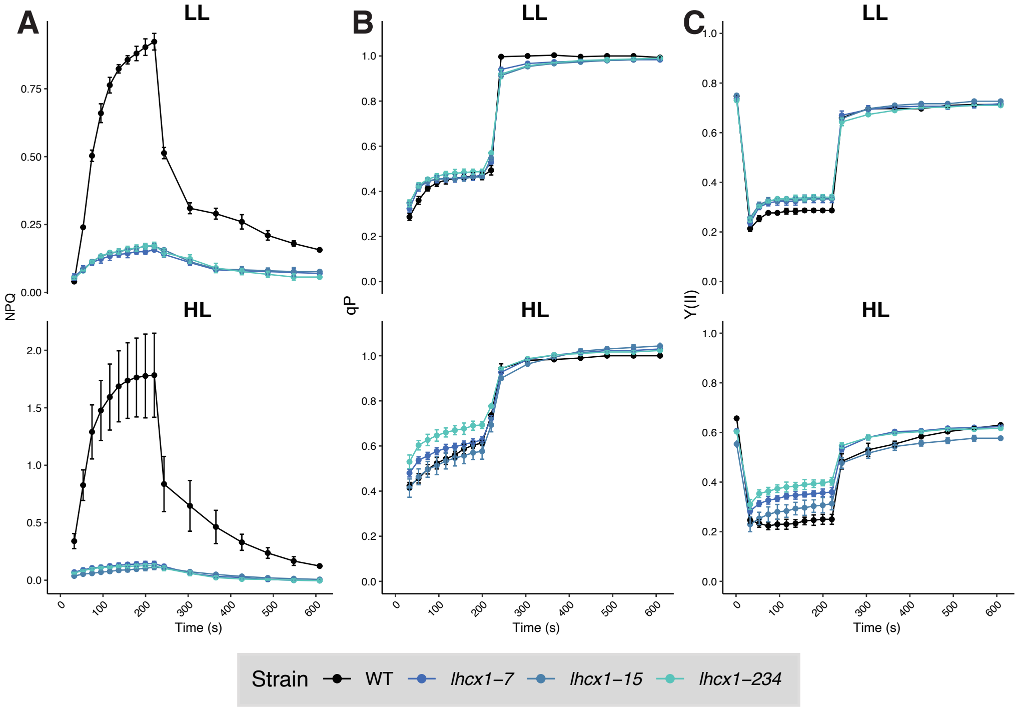


**Supplemental Figure 9. PAM Fluorescence Measurements of *C. gracilis* Wild-Type and *lhcx1* Strains Cultured with 3% $CO_2$ Aeration for three Days Under Low Light (LL) and High Light (HL)**

A) NPQ of LL and HL cultures. B) qP of LL and HL cultures. C) PSII effective quantum yield Y(II) of LL and HL cultures. The points in each figure represent mean ± standard deviation with biological replicates of $n = 3$.

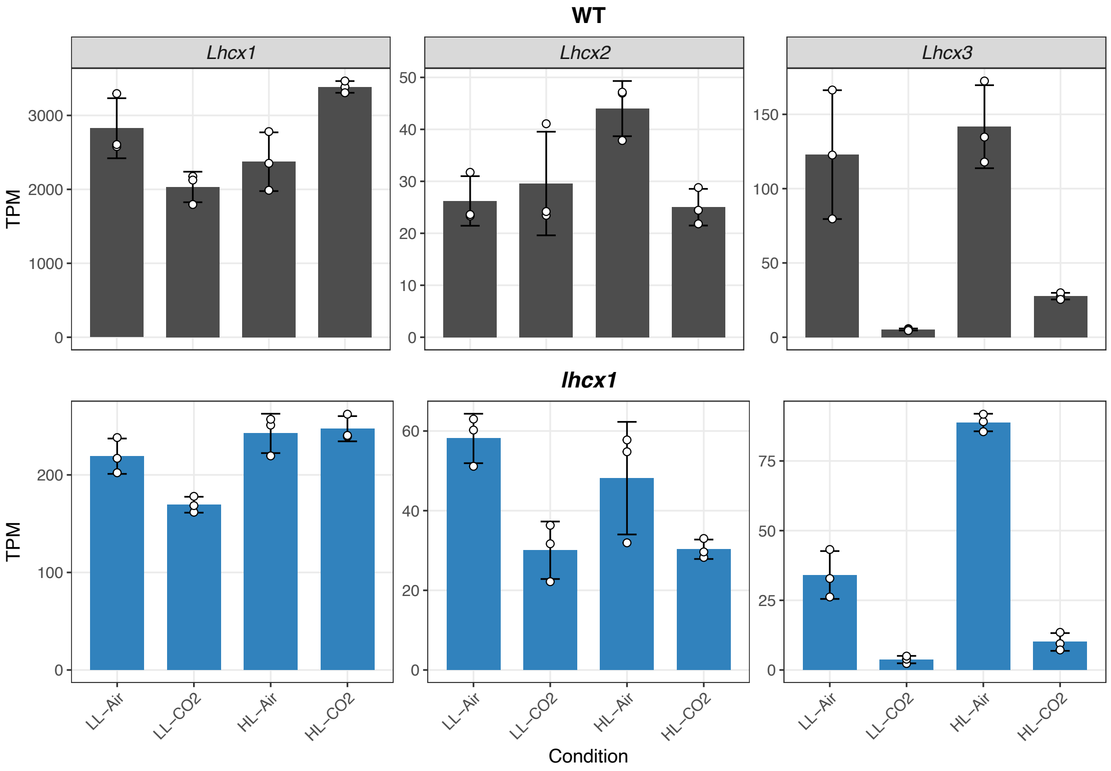


**Supplemental Figure 10. TPM of *CgLhcx1–3*.**

The means of the expression level of *CgLhcx1–3* in transcriptomes are shown as bars, and standard deviations are shown as error bars. Each replicate is indicated as a dot (n = 3, respectively).
